# An interplay between cellular growth and atypical fusion defines morphogenesis of a modular glial niche

**DOI:** 10.1101/2021.05.09.443326

**Authors:** Maria Alexandra Rujano, David Briand, Bojana Ðelić, Pauline Spéder

**Author notes:** **Corresponding author:** Pauline Spéder, PhD, Department of Developmental and Stem Cell Biology, *Brain plasticity in response to the environment* Group, Institut Pasteur/CNRS UMR3738, 25 rue du Docteur Roux, 75015 PARIS.

## Abstract

Neural stem cells (NSCs) are embedded in a multi-layered, intricate cellular microenvironment supporting their activity, the niche. Whilst shape and function are inseparable, the morphogenetic aspects of niche development are poorly understood. Here, we use the formation of the glial network of a NSC niche to investigate acquisition of architectural complexity. Cortex glia (CG) in Drosophila regulate neurogenesis and build a reticular structure around NSCs. We first show that individual CG cells grow tremendously to ensheath several NSC lineages, eventually spanning the entire tissue while partitioning the NSC population. Elaborate proliferative mechanisms convert these cells into syncytia rich in cytoplasmic bridges. Unexpectedly, CG syncytia further undergo homotypic cell-cell fusion, relying on defined molecular players of cell fusion such as cell surface receptors and actin regulators. Exchange of cellular components is however dynamic in space and time, a previously unreported unique mechanism. This atypical cell fusion remodels cellular borders, restructuring the CG syncytia. Ultimately, the coordination of growth and fusion builds the multi-level architecture of the niche, and creates a modular, spatial partition of the NSC population. Our findings provide novel insights into how a niche forms and organises while developing intimate contacts with a stem cell population.

## Introduction

Across tissues and organisms, the niche is a tailored cellular environment which/that regulates and supports stem cell behaviour by providing a structural (cell contacts and tissue topology) and signalling (biochemical cues) scaffold^1^. Despite this prominent role indissociable from stem cell activity, and hence tissue formation and homeostasis, niche cells remain poorly understood. This is particularly the case in the nervous system, where neural stem cells (NSCs) self-renew while generating new cells during neurogenesis. The NSC niche is highly complex and heterogeneous, with a diversity of cell types and interactions^2–4^ that provide extrinsic cues regulating NSC behaviour^5–8^. In mammals, neurogenic niches comprise multiple cell populations including glial cells, neurons, resident immune cells and blood vessels forming the blood-brain barrier, as well as acellular components^9–11^. The NSC niche exhibits intricate, tight cellular arrangements, such as astrocytic extensions packed in between and contacting NSCs and blood vessels^9,11^. Direct couplings also exist between several cell types, including between and within progenitor and glia populations, creating complex cellular networks sharing signals^12,13^. The NSC niche ultimately forms a functional and physical unit with specific cellular and molecular properties providing cell-cell, paracrine and systemic signals^4,14^. The niche starts to form very early during embryogenesis and becomes progressively more elaborate with the progression of neurogenesis and the acquisition of tissue complexity^11,15^. Niche composition and structure must therefore be very dynamic in order to accommodate the substantial tissue remodelling which results from neurogenesis throughout life. However, still little is known about the cellular processes involved and the supporting mechanisms happening in the niche.

In particular, we still have scarce understanding on how niche structure is established from individual cells, and how it acquires its 3D organization. Answering these questions requires being able to identify, track and manipulate independently niche cell populations *in vivo*, within their physiological context, conditions that the complexity of the mammalian brain makes challenging to achieve. First, the mammalian NSC niche has a highly heterogeneous cellular composition and architecture. In addition, mammalian models have complex genetics and the existence of multiple, parallel and tractable systems are rare. Finally, while *in vivo* models are crucial to acquire an accurate spatial and temporal picture of the cellular dynamics taking place within a 3D niche, access to a whole living brain in mammals is still difficult. To overcome these issues while offering a system allowing the investigation of core, conserved cellular and molecular mechanisms supporting NSC niche formation, we use the developing larval Drosophila brain as a model system.

Drosophila NSCs (historically called neuroblasts) are specified during embryogenesis and start proliferating to generate the neurons and glia that will form the larval CNS^16–18^. When these primary lineages are completed, embryonic NSCs exit the cell cycle and enter a quiescent state. Subsequently, during larval development, NSCs are woken up from this dormant phase^19^ by a feeding-induced nutritional signal, leading them to enlarge, re-enter the cell cycle and resume proliferation^20–23^. This second wave of neurogenesis lasts until the end of larval life, generating secondary lineages which will make up most of the adult CNS. Proliferating larval NSCs reside in a neurogenic niche which comprises common players, with related functions, to the mammalian niche –namely glial cells, a blood-brain barrier, and neurons (Figure 1a). The blood-brain barrier is essential to neurogenesis by relaying the systemic nutritional cues that will trigger NSC reactivation^22,24^. Beneath the blood-brain barrier lie the cortex glia (CG). CG display a striking structure around actively cycling NSCs, individually encasing them and their newborn progeny within membranous chambers while forming a network spanning the whole CNS (Fig. 1a-c)^25–27^. CG perform genuine niche functions. They protect NSCs against oxidative stress and nutritional restriction^28,29^, support NSC cycling^30,31^ and are essential for neuronal positioning and survival^25,27,30,32,33^. Importantly, CG network and NSC encasing are not present at the beginning of larval life, when NSCs are quiescent. Previous studies have shown that this network forms progressively in response to both nutritional cues and signals from NSCs, pinpointing an exquisite coordination between neurogenic needs, systemic cues and niche morphogenesis^27,34^.

**Figure 1:**
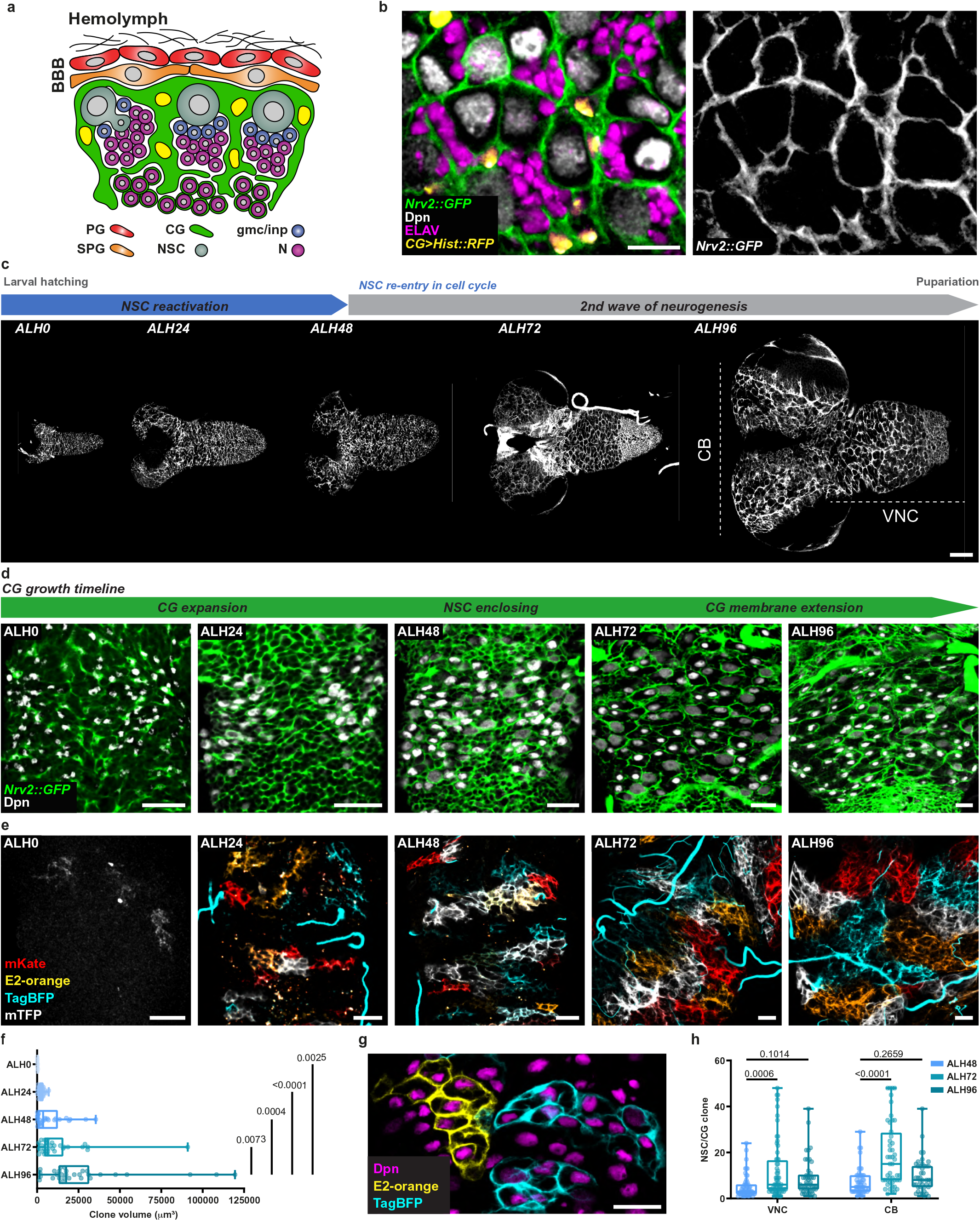
Growth of individual CG cells results in a tiled organization of the cortex glia network. a) Schematic of the Drosophila NSC niche depicting the blood brain barrier (BBB), which is made by the perineurial glia (PG, red) and subperineurial glia (SPG, orange), the cortex glia (CG, green), neural stem cells (NSC, grey), ganglion mother cells/intermediate progenitors (gmc/inp, blue) and neurons (N, magenta). b) Ventral region in the larval ventral nerve cord (VNC) at ALH72 (at 25°C) labelled with markers for the CG membranes (*Nrv2::GFP*, green), CG nuclei (*CG > Hist::RFP*, yellow), NSC (anti-Dpn, grey) and neurons (anti-ELAV, magenta). The right panel shows the CG membrane separately. Scale bar: 10 μm. c) Timeline of neurogenesis (top scheme) and assessment of CG network organization during larval development in the entire CNS at ALH0, ALH24, ALH48, ALH72 and ALH96 (at 25°C). Two main neurogenic regions are the central brain (CB), comprising two hemispheres, and the ventral nerve cord (VNC). CG membranes are labelled with *Nrv2::GFP* (ALH0, ALH72) and *CG>CD8::GFP* (ALH24, ALH48 and ALH96). Scale bar: 50 μm. d) Progressive growth and adaptation of the CG network to NSC lineages in the VNC visualized at ALH0, ALH24, ALH48, ALH72 and ALH96 (at 25°C). CG membranes are labelled with *CG>CD8::GFP* (ALH0) and *Nrv2::GFP* (ALH24, ALH48, ALH72 and ALH96). NSCs are labelled with Dpn (grey). Scale bars: 20 μm. e) Analysis of individual CG growth over time by multicolour lineage tracing using Raeppli. Images were acquired at ALH0, ALH24, ALH48, ALH72 and ALH96 (at 25°C). Constitutively expressed Cyp-Flp was used for the visualization of clones at ALH0. Hs-Flp and heat shock induction at 37°C at ALH0 was used for the visualization of clones at ALH24, ALH48, ALH72 and ALH96. Scale bars: 20 μm. f) Volume quantification of Raeppli clones in the VNC at ALH0 (n=7), ALH24 (n=25), ALH48 (n=25), ALH72 (n=32) and ALH96 (n=30). n, number of clones. Results are presented as box and whisker plots. Whiskers mark the minimum and maximum, the box includes the 25th–75th percentile, and the line in the box is the median. Individual values are superimposed. Data statistics: ordinary one-way ANOVA with a Tukey’s multiple comparison test. g) Individual TagBFP (cyan) and E2-orange (yellow) Raeppli clones encasing several NSC labelled with Dpn (magenta). Scale bar: 20 μm. h) Number of NSCs per CG clone quantification in the central brain (CB) and the VNC at ALH48 (n=53 and 51 CB and VNC, respectively), ALH72 (n=64 and 48 CB and VNC, respectively) and ALH96 (n=46 and 42 CB and VNC, respectively). n, number of clones. Bars represent the mean and the error bars are the standard deviation. Data statistics: two-way ANOVA with a Dunnett’s multiple comparison test.

Here, we used CG network morphogenesis to study niche formation and acquisition of architectural complexity. We showed that growth of individual CG cells coupled with elaborate proliferative strategies create a network of contiguous glial syncytia that ensheath subsets of NSCs. Notably, CG territories can be reshaped by an atypical cell-cell fusion mechanism, which is highly dynamic in time and space. Both CG growth and homotypic fusion are required for correct network architecture. Ultimately, we identified a niche organised in architectural units creating a spatial, modular division of the NSC population. These partitions can be remodelled by CG fusion events, resulting in a changing map of CG cells and as such NSC subsets. Importantly, the CG structure, made of connected cells capable of sharing information, and organized in spatial territories, is reminiscent of the astrocytic networks present throughout the mammalian brain^35^. Our findings provide a novel framework to understand how complex reticular structures are formed, as well as a tractable model to decipher the impact of niche structure on NSC functions and their organisation as a population.

## Results

### Growth of individual CG cells results in a tiled organization of the cortex glia network

We first sought to visualise the spatiotemporal dynamics of CG network morphogenesis during neurogenesis in the larval CNS. For this, we used either the protein trap Nrv2::GFP that labels CG membranes, or expression of membrane targeted GFP (mCD8::GFP) driven by *cyp4g15-GAL4* (expressed mostly in CG as well as in some astrocyte-like glia, readily identifiable based on morphology and dorsal compartmentalisation, see Supp. Fig S1a). In accordance with CG chambers being progressively formed in parallel with NSC reactivation^27^, the CG network starts as a loose, gaping meshwork at ALH0 (ALH: after larval hatching) that progress to a highly interconnected reticular network around ALH48, when it encloses each individual NSCs (Figure 1c-d, shown in the CNS region of the ventral nerve cord, VNC). Eventually, the CG network spans the entire tissue at ALH96. Network growth and acquisition of complexity is associated with dramatic changes in the size and morphology of CG cells, that extend their membranes to gradually accommodate the growing NSC lineages (Figure 1d). Remarkably, the resulting intricate network efficiently maintains the spatial individualities of each NSC lineage.

Next, we determined the contribution of each individual CG cell to network formation and NSC encapsulation. We expressed in CG the multicolour lineage tracing tool Raeppli^36^, that contains one single copy of membrane targeted Raeppli (Raeppli-CAAX) and can be induced at the desired time upon Flippase (FLP) recombination (Supp. Fig S1b). Its induction just After Larval Hatching (ALH0-2) resulted in the expression of exclusively one of four different colours in the CG cells. Clones extended from ALH0 to ALH96, spanning the whole tissue and forming clear boundaries between them, ultimately tiling the entire brain (Figure 1e). A similar tiled organisation was observed previously, using stochastic expression of two fluorophores, around mature neurons^32^. Quantifying the volume of individual clones over time (Figure 1f) revealed a steady growth of single colour clones from ALH0 to ALH96, with the most significant increase between ALH72 and ALH96 in concomitance with NSC lineage expansion. Remarkably, we also observed that each single CG clone (derived from one single CG cell) can encase several NSC lineages (Figure 1g), ranging from 5 NSCs per clone at ALH48 to an average of 10 NSCs per clone at ALH72 (Figure 1h). All together these results show that CG are able to grow until entirely tiling the brain while precisely encapsulating several NSC lineages.

### CG cells exhibit multiple cell cycle strategies

We then asked about the cellular mechanisms at play to support such extensive clonal growth. Two powerful, rather opposite strategies can fuel the generation of large clones. Mitosis results in both cellular and nuclear divisions and thus leads to increased cell numbers. On the other hand, endoreplication results in increased DNA content (i.e., polyploidization) without cellular division, and results in larger cell size^37–39^.

CG proliferation had been reported previously based on nuclei counts, in clones or in specific CNS region^26,32,40^. However, the cell cycle mechanisms supporting such proliferation, as well as the resulting cellular organization remained debated. While increased nuclei numbers suggested mitotic events, there were also evidence fitting endoreplicative processes, such as replication without increase in nuclei numbers detected at very early stages (ALH0-24)^34^. We thus decided to do a thorough examination of the cell cycle in CG. We first confirmed that CG nuclei numbers in the entire CNS largely increase between ALH48 and ALH96, suggesting that proliferation is enhanced when NSC lineages are expanding (Supp. Fig. S1c-d). To determine the contribution of the individual CG cells present at ALH0 to this increase, we induced Raeppli-CAAX clones at ALH0 and stained for the pan-glial marker Repo (Supp. Fig. S1e). Counting the number of Repo^+^ nuclei in each CG clone revealed a fivefold increase between ALH48 and ALH96 (Supp. Fig. S1f), in accordance with whole CNS count.

We then used the genetic tool Fly-FUCCI that allows to discriminate between G1, S and G2/M phases^41^ to assess CG cell cycling activity along network formation, focusing on the VNC for simplicity (Fig. 2a-b). FUCCI relies on a combination of fluorescently-tagged degrons from Cyclin B and E2F1 proteins which are degraded by APC/C and CRL4^CDT^^2^ from mid-mitosis and onset of S phase, respectively (Supp. Fig S2a). While CG nuclei appeared mostly in G1 at ALH0, we observed a progressive increase in the number of nuclei in S and G2/M between ALH24 and ALH72, followed by a sharp return to G1 at ALH96 (Fig. 2a-b), a temporal pattern reminiscent of the timing and level of NSC proliferation overtime. We also noticed that such change in cell cycle profile followed an antero-posterior pattern (compare ALH24 with ALH48 in Fig. 2a). This suggests that at least part of the CG population cycles between replicative and gap or mitotic phases, and that such cycling is spatially regulated and temporally coordinated with NSC behaviour.

**Figure 2:**
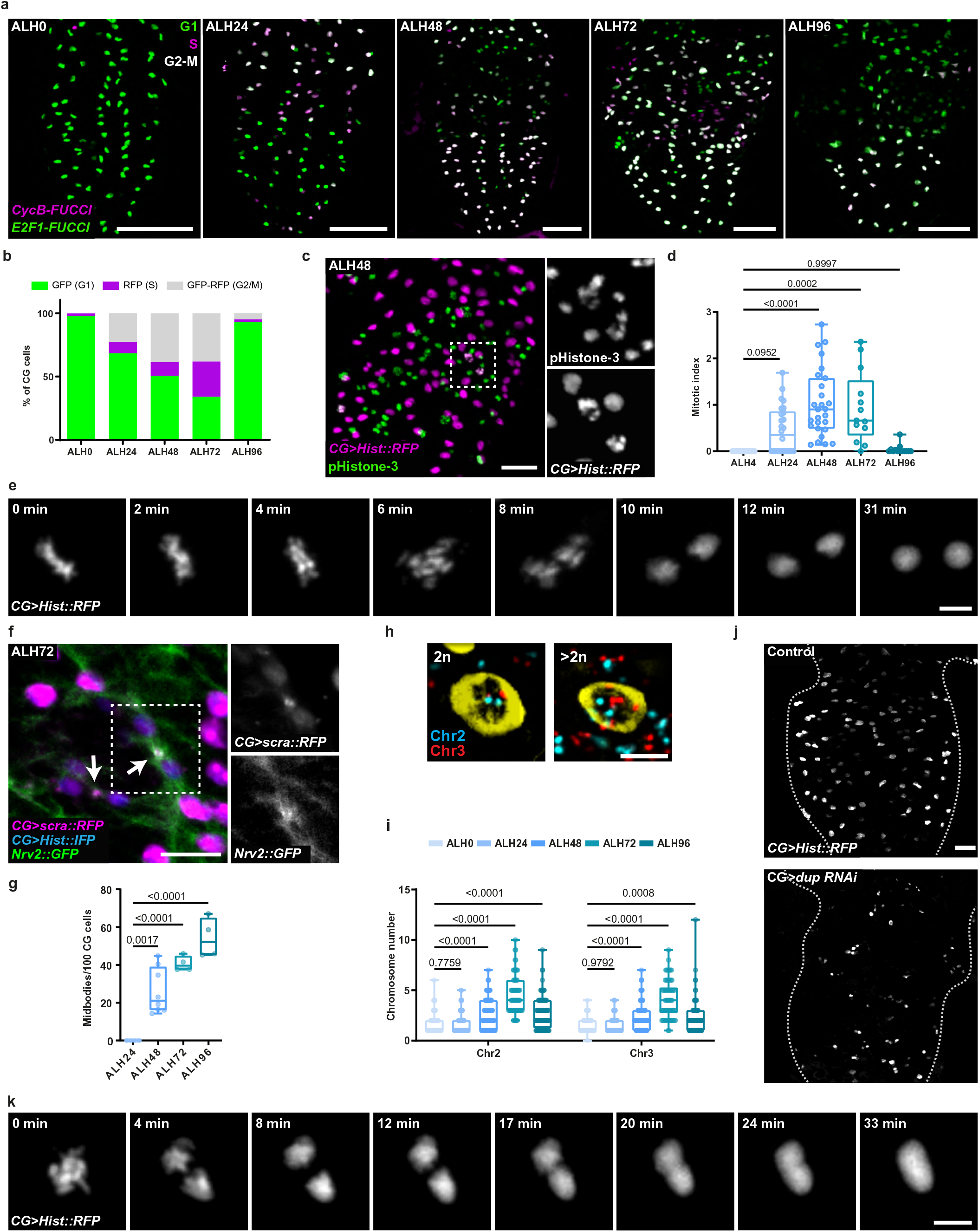
CG cells exhibit multiple cell cycle strategies. a) G1 (green), S (magenta) and G2/M (grey) phases of the cell cycle along CG network detected with Fly-FUCCI. FUCCI sensors are labelled in magenta (CycB) and green (E2F1). Scale bar: 50 μm. b) Quantification of cell cycle phase distribution in CG by Fly-FUCCI at ALH0 (n=11), ALH24 (n=15), ALH48 (n=23), ALH72 (n=13) and ALH96 (n=6) (at 25°C). n, number of CNS analysed. Stacked bars represent the percentage of cells in each phase. c) Representative image of a larval VNC expressing Hist::RFP in CG (magenta) and stained with phospho-histone H3 antibody (pHistone-3, green) to visualise mitotic CG nuclei (grey). Scale bar: 20 μm. Higher magnification of separate channels from the region inside the dashed rectangle are shown on the right. d) CG mitotic index quantification in larval CNS at ALH0 (n=15), ALH24 (n=26), ALH48 (n=27), ALH72 (n=13) and ALH96 (n=13) (at 25°C). n, number of CNS analysed. Results are presented as box and whisker plots. Whiskers mark the minimum and maximum, the box includes the 25th–75th percentile, and the line in the box is the median. Individual values are superimposed. Data statistics: ordinary one-way ANOVA with a Tukey’s multiple comparison test. e) Still images of a time-lapse movie (Movie S1) of mitotic CG expressing *Hist::RFP* (grey). Scale bar: 5 μm. f) Expression of *mRFP::scra* (magenta) in CG to monitor contractile ring and midbody formation. CG membranes and nuclei are labelled with *Nrv2::GFP* (green) and *Hist::IFP* (blue) respectively. Arrows indicate midbodies/contractile ring. Scale bar: 10 μm. Higher magnifications of *mRFP::scra* and *Nrv2::GFP* separate channels from the region demarcated by the dashed rectangle are shown on the right. g) Quantification of the number of midbodies per 100 CG cells in larval VNCs at ALH24 (n=4), ALH48 (n=8), ALH72 (n=4) and ALH96 (n=4) (at 25°C). n, number of VNCs analysed. Results are presented as box and whisker plots. Whiskers mark the minimum and maximum, the box includes the 25th–75th percentile, and the line in the box is the median. Individual values are superimposed. Data statistics: ordinary one-way ANOVA with a Tukey’s multiple comparison test. h) Fluorescence in situ hybridization (FISH) using probes for chromosomes 2 (Chr2, cyan) and 3 (Chr3, red) in CNS expressing nls::LacZ (yellow) to mark the CG nuclei. 2n (upper) and >2n (bottom) nuclei are shown. Scale bar: 5 μm. i) Quantification of FISH signals in CG nuclei at ALH0 (n=95), ALH24 (n=189), ALH48 (n=140), ALH72 (n=70) and ALH96 (n=108). N, number of CG cells analysed. Results are presented as box and whisker plots. Whiskers mark the minimum and maximum, the box includes the 25th–75th percentile, and the line in the box is the median. Individual values are superimposed. Data statistics: two-way ANOVA with a Dunnett’s multiple comparison test. j, k) CG nuclei (j, *CG > Hist::RFP*) and CG network (k, *Nrv2::GFP*) in control CNS and in CNS where CG-specific downregulation of doubled-parked (*dup RNAi*) was induced. Scale bar: 20 μm. l) Still images of a time-lapse movie (Movie S3) of a CG expressing *Hist::RFP* (grey) undergoing endomitosis. Scale bar: 5 μm.

To assess whether CG cells undergo proper mitosis, we checked *bona fide* mitotic hallmarks. We first stained CG cells with the mitotic marker phospho-histone H3 (PH3, Fig. 2c-d) and detected PH3^+^ CG cells between ALH24 and ALH72, fitting the FUCCI window with more CG cells in S or G2/M phases. Next, by performing live-imaging of RFP-tagged histone (*Hist::RFP*) driven in CG on whole brain explants (see Methods), we were able to observe DNA condensation, metaphase alignment and chromosomes’ segregation (Fig. 2e, Movie S1). Moreover, we observed nuclear envelope breakdown followed by reformation using *Lamin::GFP* expressed in CG (Supp. Fig. S2b, Movie S2). We also looked at the behaviour of the Drosophila homolog of anillin (*scraps, scra*), a conserved scaffolding protein involved in late stages of cytokinesis^42^. Anillin is found in the nucleus during interphase and relocates to the contractile ring during cytokinesis ^43^. It then forms part of the midbody, a contractile ring-derived microtubule-rich proteinaceous structure assembled at the intercellular bridge between the two daughter cells at the end of mitosis and that marks the abscission site. Expressing RFP-tagged anillin in CG (*mRFP::scra)* uncovered midbody-like structures in between recently divided CG (Fig. 2f, identified by a decrease in nuclear-localised anillin compared to neighbouring CG nuclei) and along the CG membranes (Supp. Fig S2c). Quantifying anillin-positive midbody structures along time (Fig. 2g) revealed an increase between ALH48 and ALH96, paralleling FUCCI and PH3 windows. All together, these data suggest that CG cells do undergo proper mitosis, including nuclear division and cytokinesis up to midbody formation.

Next, we sought to address whether endoreplication and subsequent polyploidization could also happen in CG. We assessed CG ploidy through DNA Fluorescence *In Situ* Hybridization (FISH) on chromosomes II and III (two out of the four Drosophila chromosomes^44^) in labelled CG nuclei along development (Figure 2h-i). We observed that at early stages, CG have a normal ploidy of 2n, which increases at ALH72 for part of the CG population, and decreases again to 2n at ALH96. Although we cannot exclude that part of this increase corresponds to catching DNA replication before mitosis (PH3^+^ staining also peaks at ALH48-72, Fig. 2d), odd numbers as well as n>4 imply a contribution of polyploidization. Moreover, CG-specific downregulation of Dup (*double parked* gene), a DNA replication protein shown to be crucial for endoreplication^45,46^, caused a strong reduction in CG nuclei size and number (Figure 2j). Notably, endoreplication covers two cell cycle variants^37,47^. Endocycle alternate DNA replication (S-phase) with a gap (G) phase and do not show any mitotic features. Endomitosis includes S phase and some aspects of mitosis up to telophase^48^, but do not complete cellular division. By live-imaging on whole brain explants, we were able to observe endomitotic events, characterized by entry into mitosis followed by chromosomes segregation but absent later mitotic stages, instead with the DNA collapsing back into only one nucleus (Figure 2k, Movie S3). All together, these data show that polyploidization does occur in CG in a temporary fashion, in some cases through endomitosis.

### CG glia are syncytial units

While CG displayed well-characterized marks covering different mitotic steps, we also noticed peculiar behaviours that indicated a subtler picture. First, using live-imaging, we noticed that mitoses often appeared synchronised between several nuclei (Fig. 3a, Movie S4). Similarly, using Fly-FUCCI, groups of neighbouring nuclei were found at the same cell cycle phase (Fig. 3b). Moreover, we observed that several close-by CG nuclei were undergoing cytokinesis at the same time, even sometimes seemingly linked by anillin cytoplasmic staining (Fig. 3b). Such coordinated behaviour between a group of CG nuclei suggest that they are receiving the same cell cycle cues. We thus wondered whether multiple CG nuclei could actually be sharing cytoplasmic material.

**Figure 3:**
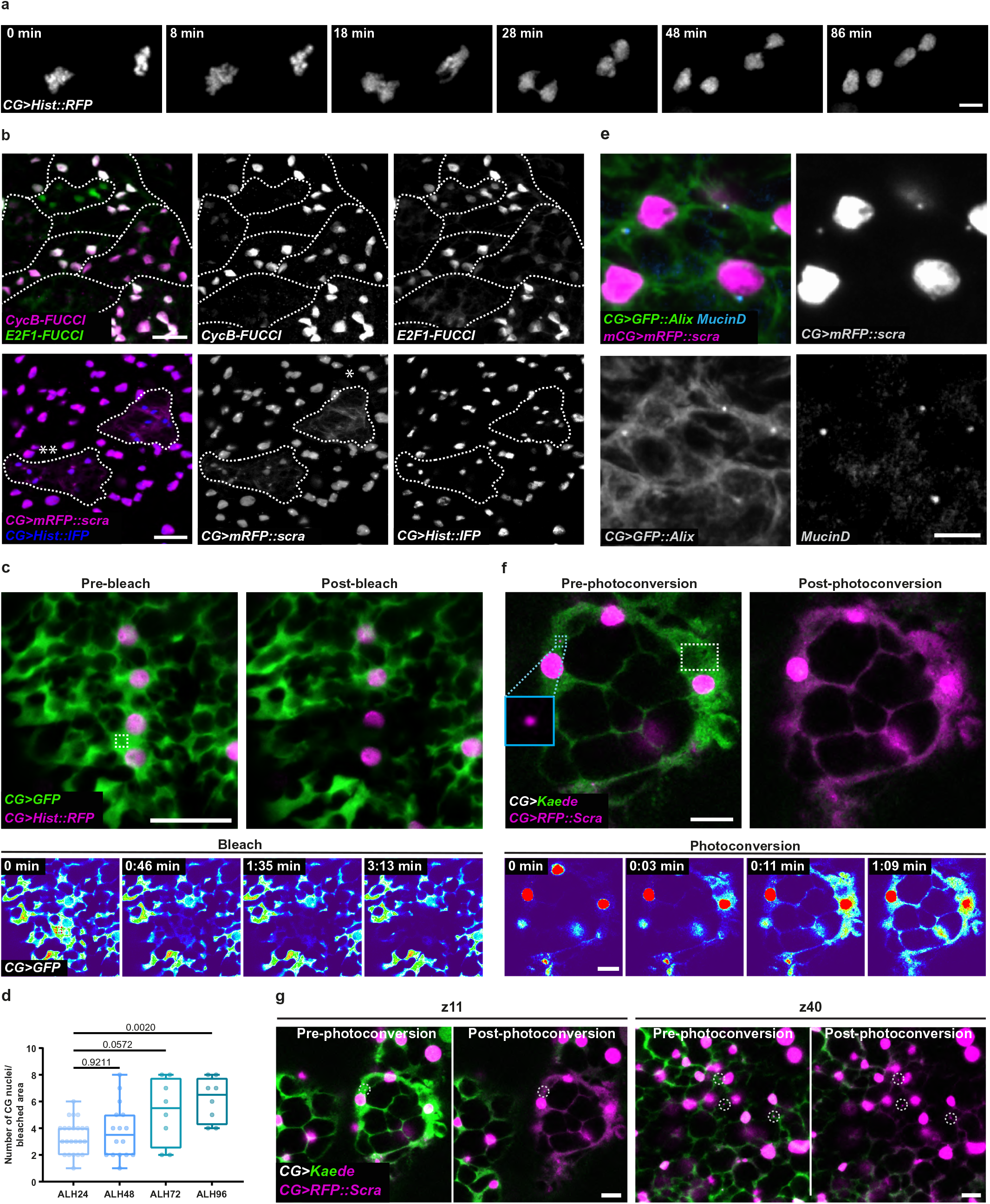
CG glia are syncytial units. a) Still images of a time-lapse movie (Movie S4) of two CG expressing *Hist::RFP* (grey) undergoing mitosis synchronously. Scale bar: 5 μm. b) Synchronous behaviour of CG observed with Fly-FUCCI (left panels), where clusters of CG are found at the same cell cycle phase, and with anillin staining that also show clusters of CG undergoing mitosis (*) and cytokinesis (**) at the same time (right panels). Synchronous clusters are delineated with dashed lines. FUCCI sensors are labelled in magenta (CycB) and green (E2F1). Anillin is labelled with mRFP::scra (magenta) and CG nuclei with Hist::IFP (blue). Separate channels are shown in the bottom. Scale bars: 20 μm. c) Sharing of cytoplasmic material between CG assessed by Fluorescence Loss In Photobleaching (FLIP) of cytosolic GFP (green). Top panels depict a region in the VNC before (pre-bleach) and after bleaching (post-bleach). CG nuclei are labelled with *Hist::RFP* (magenta). Bottom panels show intermediate time points (GFP only, pseudocolored with thermal LUT) during continuous photobleaching. Bleached area is delineated by the dashed square. Scale bars: 20 μm. d) Quantification of the number of CG nuclei in the bleached region after FLIP at ALH24 (n=23), ALH48 (n=16), ALH72 (n=8) and ALH96 (n=8). n, number of FLIP experiments analysed. Results are presented as box and whisker plots. Whiskers mark the minimum and maximum, the box includes the 25th–75th percentile, and the line in the box is the median. Individual values are superimposed. Data statistics: two-way ANOVA with a Dunnett’s multiple comparison test. e) Anillin (mRFP::scra, magenta) is found in punctated structures enriched in Alix (*CG>UAS-GFP::Alix*, green) and Mucin-D (anti-Mucin-D, cyan), two known components of midbodies and intercellular bridges. CG nuclei are stained with His::IFP (*CG>His::IFP*, grey). Scale bar: 5 µm. f) CG connection via the midbodies marked by anillin (mRFP::scra, magenta) assessed by photoconversion of cytosolic Kaede from GFP (green) to RFP (magenta, cKaede). Top panels depict a region in the VNC before (pre-photoconversion) and after photoconversion (post-photoconversion). The photoconverted area is delineated by the white dashed square and an isolated midbody (clear blue inset) is shown in between CG cells. Bottom panels show intermediate time points (RFP/cKaede only, pseudocolored with thermal LUT) during continuous photoconversion. Scale bars: 10 μm. g) Selected slices (z11, left panel and z40, right panel) from Z-stacks before and after photoconversion shown in f) are displayed next to each other. The cKaede signal overlaps with several midbodies-like (mRFP::scra puncta, dashed white circles) throughout the Z-stack, showing that CG units are rich in intercellular bridges. Scale bar: 10 µm. z-step is 0.50 µm.

To test this, we relied on a Fluorescence Loss In Photobleaching (FLIP) technique, an approach used to examine the movement of molecules inside a cell and that can also serve to assess the continuity of a cellular compartment (reviewed in ^49,50^). FLIP relies on the continuous bleaching of a small region of a fluorescently-labelled cell, while recording the entire zone whose continuity is being assessed. The continuous illumination will result not only in the bleach of the targeted region, but also will lead to the loss of fluorescence in any connected area, due to molecular diffusion. In contrast, non-connected areas will not be bleached. We expressed cytoplasmic GFP and RFP-tagged histone (*Hist::RFP*) in the entire CG population and imaged an area containing several CG nuclei. We then repetitively bleached GFP in a small region of the cytoplasm and recorded the loss of fluorescence with respect to CG nuclei. Strikingly, we were able to observe loss of fluorescence in large areas containing several CG nuclei (Fig. 3c and Supp. Fig. S3a), implying that indeed these CG nuclei are part of a continuous, connected cytoplasmic compartment. Quantifying FLIP experiments at different times revealed that the average number of connected CG nuclei increases twofold along CG network formation (Fig. 3d; average ALH24 = 3, versus average ALH96 = 7). These experiments show that CG cells are thus multinucleated.

Endomitosis could produce multinucleated cells in the rarer case they go through nuclear envelope breakdown and reformation. Nevertheless, a straightforward explanation to account for such an extent of multinucleation would be that CG undergo mitosis but fail to complete cytokinesis. The midbody is indeed a temporary structure formed between the two daughter cells during cytokinesis. While recent studies have shown that midbodies can be retained and have roles beyond cytokinetic events^51^, their main function is linked to abscission, after which their usual fate is to be cleaved and discarded. In some instances though, the midbody can be conserved at the site of cleavage to become a stable cytoplasmic bridge keeping the two daughter cells connected^52,53^. In this case, the midbody grows and matures to become a ring-type structure (often coined ring canal) allowing exchange of large molecules.

Interestingly, counting anillin-enriched midbody structures along CG membranes revealed a steady increase in numbers over time, including up to the end of larval stage (Fig. 2g), what entails that they are not discarded but rather remain. This suggests that CG cells enter mitosis but at least in some cases, fail cytokinesis, staying connected by the midbody and related intercellular bridge. We then wondered whether other proteins known to associate with the midbody and ring canals were also present in puncta along CG membranes. We first found that a GFP fusion of ALIX, an ESCRT-associated scaffold protein required for abscission in the fly germline^54^, and endogenous Mucin-D, a mucin-type glycoprotein identified as a generic component of Drosophila intercellular bridges^55^, were co-localising with or adjacent to mRFP::scra puncta along CG membranes, respectively (Fig. 3e and Supp. Fig. S3b). In addition, Mucin-D puncta co-stained for a fluorescent fusion of the kinesin-like Pavarotti, an essential component of the contractile ring and derived structures^56^ (Supp. Fig. S3c). Mucin-D and GFP::Pavarotti puncta exist independently of mScra::RFP expression, indicating that anillin overexpression does not induce their artificial recruitment. These data indicate that midbody-type structures containing multiple molecular components of midbody and stable intercellular bridge are present along CG membranes.

To demonstrate that CG cells stay connected through such structures, we first performed FLIP, expressing a cytoplasmic GFP together with mRFP::scra in all CG cells (Fig. S3d). We repetitively bleached GFP in a small cytoplasmic region next to an isolated Scra^+^ punctum localised in a narrow cytoplasmic extension between CG nuclei. We found that the loss of fluorescence was able to propagate through the Scra^+^ punctum, reaching CG nuclei localised on the other side. To fully demonstrate the existence as well as extent of such cytoplasmic connection, we then expressed in the CG, in combination with mRFP::scra, a photoconvertible protein (Kaede^57^), that irreversibly switches from GFP to RFP when excited by UV pulses. Excitation of a small CG zone led to the propagation of the converted form (herein named cKaede) in the whole plane, including through a Scra^+^ punctum, ultimately covering the latter own fluorescence and reaching several CG nuclei (Fig. 3f). Z-imaging of the cKaede signal before and after localised photoconversion revealed a connected zone covering several nuclei and Scra^+^ puncta (Fig. 3g and Movie S5). All together, these data show that CG cells are multinucleated and form syncytial compartments throughout which cytoplasmic proteins can shuttle, and which result in part from incomplete cytokinesis that leave cells connected via the midbody/intercellular bridge. From now on, we will call these syncytial structures, originating from mononucleated cells and encapsulating several NSCs (Fig. 1g-h), CG units.

### CG units can undergo cellular fusion

Multicolour clonal analysis with membrane targeted Raeppli showed that individual CG cells give rise to neighbouring units with well-defined boundaries that tile the CNS (Fig. 1e, g). Intriguingly, we were also able to observe membrane areas with colours overlap (Fig. 4a), with numbers fluctuating over time (Supp. Fig. S4a-b). The partial nature of the overlap, as well as colour induction well before polyploidization (see Fig. 2i), excluded that such event would come from polyploid cells harbouring multiple copies of the genetic tool. We wondered whether colour sharing between two neighbouring units could be a result of cell-cell fusion. Cellular fusion is the process by which two cells merge their membranes into a single bilayer, resulting in the exchange of their cytoplasmic content and subcellular compartments. It is a key recurring event in life, from egg fertilization to organogenesis, through viral infection. Cell fusion is a stepwise operation (reviewed in^58–61^). First cells become competent for fusion, usually with one donor and one acceptor. They then adhere to each other through cell recognition molecules. Membrane hemifusion proceeds ultimately leading to pore formation. Cells start to exchange their cytoplasmic content through the pore, which widens, and eventually fully integrate, sharing all their compartments.

**Figure 4:**
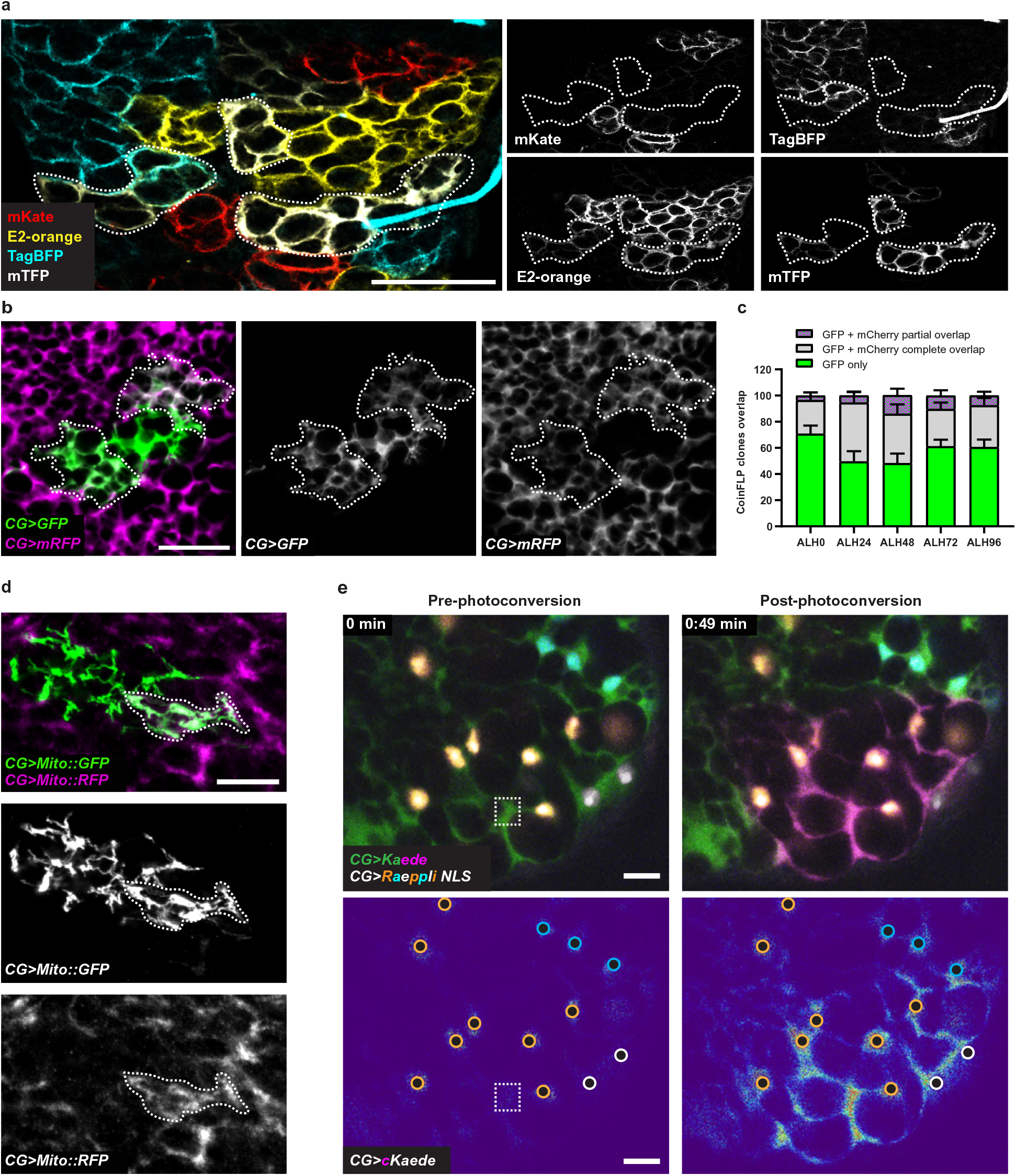
CG units can undergo cellular fusion. a) Restricted areas of colour overlapping in membrane targeted CG Raeppli clones at ALH72 (dashed lines). Scale bar: 50 μm. b) Cytoplasmic exchange between CG units assessed in CoinFLP clones (methods and Supp. Fig. S4c). Clones expressing either cytosolic GFP (green) or cytosolic RFP (magenta), show regions of partial overlapping (dashed lines). Scale bar: 20 μm. c) Quantification of areas of partial (grey-hashed green), total (grey) or no overlap (green) between clones expressing cytosolic GFP and RFP. Due to the bias in the CoinFLP system that generates very large connected clones in one colour (RFP in our case) and small sparse clones in the other colour (GFP), only green clones were taken in account for the no overlap category. Stacked bars represent the mean and error bars represent the SEM. Data statistics: two-way ANOVA with a Šídák’s multiple comparisons test. No statistically significative differences were found. d) Mitochondrial exchange between CG units assessed in Coin-FLP clones (methods and Supp. Fig. S4c). Clones expressing mitochondrial markers Mito::GFP (green) or Mito::RFP (magenta), show regions of partial overlap (dashed lines). Scale bar: 10 µm. e) Continuity between CG units due to cellular fusion was assessed by photoconversion of cytosolic Kaede expressed in the CG in combination with early induction of multicolour labelling of CG nuclei (Raeppli-NLS) that leads to clonal labelling of the nuclei in CG units. Iterative photoconversion was performed in a small area (dashed rectangle) within a Raeppli-NLS CG clone containing nuclei of one colour. Top panels depict the assessed area before (pre-photoconversion) and after photoconversion (post-photoconversion). Bottom panels show the converted form (cKaede) only, pseudocolored with thermal LUT) before and after photoconversion, with nuclei represented by black discs outlined with the respective Raeppli colour. In total, three different colours of nuclei are joined by the cKaede signal. Scale bars: 10 μm.

We first asked whether such partial colour overlap between clones could be detected for cellular compartments other than the plasma membrane. We took advantage of the CoinFLP technique, which allows the stochastic labelling in two colours of individual cells within the same population (Supp. Figure S4c)^62^. A bias in the system results in the generation of a minority of well-sparse clones in one of the two colours, making them easy to localise and quantify. Early induction of this tool in CG cells using *cyp4g15-FLP* (which is active before ALH0) and two differently-labelled fluorescent cytoplasmic markers (GFP and mCherry), generated three situations (Fig. 4b-c): i) a majority of clones of only one colour (GFP only, green); ii) clones fully colocalising with the other colour (GFP + mCherry complete overlap, grey); and iii) a minority of clones partially colocalising with the other colour (GFP + mCherry partial overlap, grey-hashed green). While full overlaps might come from polyploidy, at least in part, the occurrence of partial cytoplasmic overlaps fitted the hypothesis of fusion between CG units. We then performed a similar experiment this time using fluorescently-labelled mitochondrial markers, and also found partial colocalisation in some cases (Fig. 4d), suggesting that two CG units from different origins can share these organelles. Finally, we used a nuclear-tagged version of Raeppli (Raeppli-NLS) to identify the nuclei belonging to different CG units. While we observed clones of neighbouring nuclei with a tiled organisation reminiscent of Raeppli-CAAX, and confirming clonal expansion of individual CG cells (Supp. Fig. S4d), we also found intriguing overlaps at the border of clones, with few nuclei exhibiting two colours showing qualitative inverse intensities (Suppl. Fig. S4e). This suggests that nuclei from different CG units in close vicinity can exchange nuclear targeted proteins. All together, these data show that CG units can share subcellular compartments, including plasma membrane, cytoplasm, mitochondria and nucleoplasm.

A first prediction arising from the occurrence of cellular fusion between CG units would be the creation of cellular compartments (*i.e.*, with a continuity of information) containing nuclei from different origins. To test this hypothesis, we expressed Kaede in the whole CG population, together with stochastic multicolour nuclear labelling (Raeppli-NLS) induced early, hence leading to differently labelled clonal CG units (such as seen in Supp. Fig 4d). Localised photoconversion led to a signal (cKaede) that propagated from within the targeted CG clone to nuclei of other colours, belonging to adjacent CG neighbours, both in the same plane and throughout the depth of the tissue (Fig. 4e and Movie S6). We observed this event in a number of CG units with diverse organisation (see Supp. Fig. 4f for another example), making the observation reproducible qualitatively while difficult to assess quantitatively in terms of extent of connection. A similar conclusion was reached through FLIP on CG expressing a cytoplasmic fluorescent marker (GFP) in Raeppli-NLS CG clones. Continuous bleaching in a small area of the cytoplasmic GFP surrounding one of the CG clones indeed resulted in a loss of fluorescence not only in the targeted CG clone, but also in its adjacent neighbour (Supp. Fig. 4g). From these data, we can conclude that CG units can fuse in a homotypic manner, and generate connected areas from different origins, leading to exchange of subcellular compartments and associated signals at larger spatial scale.

### Fusion of CG units is dynamic and can create novel cellular compartments

Cell fusion entices that information could propagate from one cell to the other up to the end of the fused area. In classical models, the two partners fully integrate, generating one bigger cell. However, the existence of partial overlaps of cellular compartments (membrane, Fig. 4a; cytoplasm, Fig. 4b) is unusual and implies that the fusion did not lead to complete integration and sharing of all compartments between CG partners.

To determine the extent of compartmental continuity and signal propagation between the fusing/fused CG units, we combined the identification of zones of partial cytoplasmic overlap through CoinFLP (see Fig. 4b; GFP and mCherry) with a FLIP approach in live-imaging. We choose as example (n = 3) a GFP clone displaying a partial overlap with mCherry, as well as sharp borders with mCherry-only regions (Fig. 5a-e and Supp. Fig. 5-1a). Interestingly, the overlapping area also presented two sub-zones, distinguished by the GFP level (H_GFP_, high and L_GFP_, low on Supp. Fig. 5-1a). To assess the effect of the FLIP, significance of the percentage of fluorescence loss (i.e., attribution to the FLIP rather than chance) was determined on the sample itself (see Methods and Supp. Fig. 5-1a-b) and 19.1%/20.8% were identified as thresholds for GFP and mCherry respectively (confidence level 95%).

**Figure 5:**
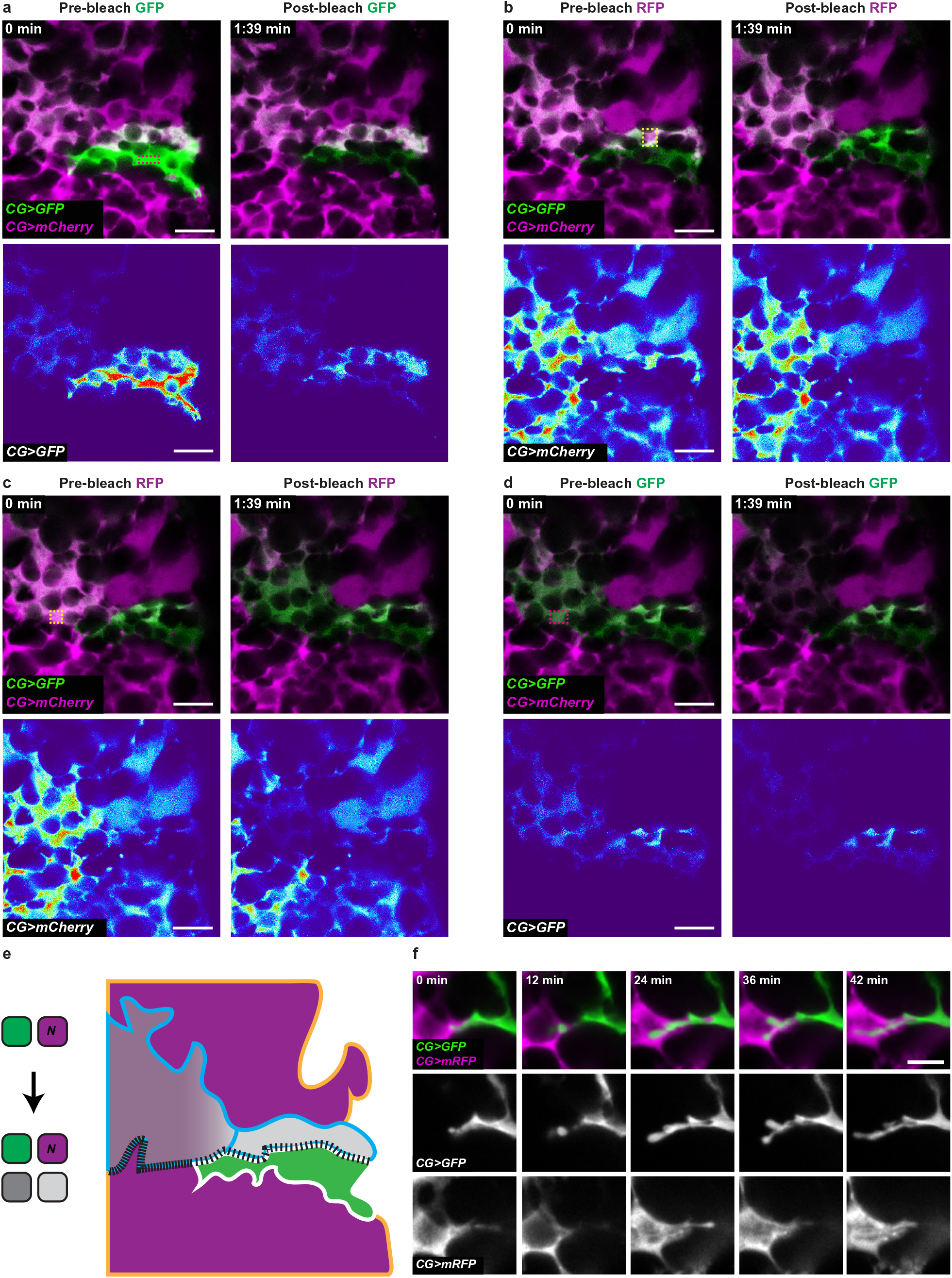
Cell fusion between CG units is atypical. a-e) Propagation of information/signals between fused areas was assessed by FLIP in clones generated by CoinFLP with cytosolic GFP (green) and RFP (magenta) in CG. A GFP expressing clone with areas of partial and no overlap with an RFP expressing clone was selected. For each experiment, continuous bleaching was performed in a small area (dashed rectangle, pink for GFP and yellow for mCherry), and loss of fluorescence in different regions was measured (see Supp. Fig. S5-1c for values). Top panels depict the assessed area before (pre-bleach) and after bleaching (post-bleach). Bottom panels show the bleached channel only, pseudocolored with thermal LUT. Scale bars: 10 μm. a) Continuous bleaching of GFP in the non-overlapping part of the GFP clone (GFP zone). b) Continuous bleaching of mCherry in the overlapping part of the GFP clone with high GFP intensities (H_GFP_ subzone). c) Continuous bleaching of mCherry in the overlapping part of the GFP clone with lower GFP intensities (L_GFP_ subzone). d) Continuous bleaching of GFP overlapping part of the GFP clone with lower GFP intensities (L_GFP_ subzone). e) Schematics of the findings from 5a-d. Left panel represents the outcome of fusion between CG units from experiments 5a-d, with N describing the unknown number of original mCherry cells. As we cannot know the number of mCherry original cells in the movie area, we cannot distinguish between whether i) the GFP clone fused with two mCherry cells to independently generate L_GFP_ and H_GFP_; or ii) the GFP clone fused with one mCherry cell (mCherry 2), forming a mixed compartment that further splits. Right panel illustrates the different zones distinguished by the overlap between the GFP and the mCherry signals, as well as the diffusion barriers existing at the time of recording. Solid lines indicate full diffusion barriers and dashed lines the existence of compartmental continuity, the extent of which is indicated by the interval between the dots. f) Still images of a time-lapse movie (Movie S11) of the region of interaction between two neighbouring CG clones generated with CoinFLP and expressing either cytosolic GFP or RFP, at ALH48. Scale bar: 5 μm.

First, we found that continuous localised bleaching of the GFP signal in a region devoided of any mCherry signal (GFP zone, Fig. 5a, Supp. Fig. 5-1, Supp. Fig. 5-2a and Movie S7) led to a loss of fluorescence not only in the CG clone targeted by the bleaching (≈ 87% loss), but also in the overlapping adjacent area (GFP + mCherry, zone H_GFP_) (≈ 47% loss), up to the border with another clone (mCherry alone). This shows that the overlapping zone between two CG clones is indeed in cytoplasmic continuity with at least one of them, and corresponds to area of some signal exchange. However, continuous localised bleaching of mCherry (≈ 78% loss) in the same overlapping subzone (H_GFP_) did not lead to a significant decrease of fluorescence in the adjacent mCherry region (mCherry 1; ≈ 13% loss), even when restricting our analysis to a smaller, adjacent portion (mCherry 1 small, ≈ 19% loss; Fig. 5b, Supp. Fig. 5-1, Supp. Fig. 5-2b and Movie S8). This suggests that the fused mCherry-GFP compartment does not communicate, or at least in a detectable manner, with an original mCherry^+^ clone. Moreover, we noticed that such bleaching of mCherry in the H_GFP_ zone also did not result in a significant loss in the overlapping subzone with lower GFP signal (L_GFP_, ≈ 0.4% loss). This entails that some diffusion barrier exists between the two H_GFP_ and L_GFP_ subzones. To confirm this observation, we performed the reciprocal FLIP experiment, and bleached a small area of mCherry signal in L_GFP_ (Fig. 5c, Supp. Fig. 5-1, Supp. Fig. 5-2c and Movie S9). While it led to a dramatic loss of mCherry signal in the targeted L_GFP_ subzone (70% loss), it did not affect the mCherry signal left in the H_GFP_ zone (≈ 8% loss). This again revealed a sharp diffusion barrier between the L_GFP_ and H_GFP_ subzones. However, we detected a restricted, albeit significant, decrease (≈ 22% loss) in the mCherry signal adjacent to L_GFP_ (mCherry 2 small, Supp. Fig. 5-1a). This suggests that some cytoplasmic exchange is still happening between a fused zone (L_GFP_) and a mCherry-only zone (mCherry 2), which the fused zone is likely derived from. Finally, the existence of a diffusion barrier between the L_GFP_ and H_GFP_ subzones was further confirmed by bleaching in L_GFP_ the GFP signal (Fig. 5d, Supp. Fig. 5-1, Supp. Fig. 5-2d and Movie S10), whose fluorescence loss (≈ 64% loss) did not propagate to the H_GFP_ compartment (≈ 9% loss). Surprisingly, we did find however that it led to a decrease in the GFP signal of the GFP-only area (≈ 31% loss), implying that the original GFP clone is also still connected, to some level, to L_GFP_, in addition to H_GFP_ (Fig. 5a).

All together, these data confirm that cellular fusion between CG units both happens and is atypical by its partial, dynamic nature. While it results in compartmental exchange between CG units, such sharing can be temporary, being severed or at least restricted after some time, as indicated by a remaining GFP-only compartment and a FLIP which propagates with sharp changes (*e.g.*, between L_GFP_, 70%, and mCherry 2, 22%). As such, it results in the creation of novel CG cells/units, owning features of both original CG partners, from which they can eventually separate to form compartments with their own properties (Fig. 5e).

### Cell fusion between CG units is regulated by canonical fusion molecules

A biological model which has been highly instrumental in deciphering fusion hallmarks is the generation of myofibers in Drosophila (reviewed in^63–65^). It follows a typical sequence of events (Supp. Fig. S6a): binding of the two partners, cascade of intracellular signalling, remodelling of the actin cytoskeleton and membrane hemifusion followed by pore formation. The end point is the creation of a multinucleated cell, the muscle fiber. These processes rely on key cell recognition and adhesion molecules (immunoglobulin-domain receptors: Sns, Hbs; Kirre/Duf and Rst), on adapter proteins (Rols7/Ants; Dock), as well as on the combined actions of multiple actin regulators (WASp, Rac, Scar, Arp2/3 to name a few). Adhesion through cell surface receptors and cytoskeletal remodelling are also core steps in myoblast fusion in vertebrates, involving some conserved molecular players^63,66^. Considering the atypical nature of cell fusion between CG units, we wondered whether similar molecular players, and as such, cellular events, were involved in this process.

First, using live-imaging, we assessed whether we could observe dynamic cellular behaviour at the border between CG units. Using two differently-labelled fluorescent cytoplasmic markers in a CoinFLP set up, we indeed observed active protrusion-like structures tunnelling into the reciprocal cells (Fig. 5f and Movie S11). This suggests that some cellular remodelling takes place at the interface between two fusing CG units. In addition, driving β-actin fused to CFP in CG revealed localised zones of higher activity (Supp. Fig. S5-2e). Next, we asked whether known molecular players of myoblast fusion were expressed and required for fusion between CG units. In light of the restricted and for now spatially unpredictable occurrence of fusion events in the CG, we decided to first focus on molecular players known to be expressed in the two partners. We turned to Myoblast City (*mbc*), a Guanine nucleotide Exchange Factor (GEF) implicated in actin remodelling and known to be expressed, if not required^67^, in both fusing cells (Supp Fig. S6a). We first took advantage of a genomic trap line inserting a GAL4 driver under the control of *mbc* enhancers (Trojan *mbc-GAL4*^68,69^). Driving both a nuclear (Hist::RFP) and membrane reporter (mCD8::GFP) revealed a strong expression in the CG (co-stained with the glial marker Repo), reproducing the characteristic CG meshwork pattern (Fig. 6a). Moreover, expressing lineage tracing tools (i-TRACE^70^ and G-TRACE^71^) under *mbc-GAL4* indicated that *mbc* is expressed in the CG throughout development (Supp. Fig. S6b-d). *mbc* expression in the CG was further confirmed by immunostaining with an anti-Mbc antibody, whose staining was enriched along the CG membranes (Fig. 6b), and lost upon *mbc* RNAi-mediated downregulation in the CG (Fig. 6b). In addition, we were able to detect a faint and more restricted staining for the adhesion molecule Kirre, which colocalised with a marker for the CG membrane and which was lost under *kirre* knockdown in the CG (Fig. 6c). Altogether, Mbc and Kirre, two known regulators of myoblast fusion, are expressed in the CG during larval stages.

**Figure 6:**
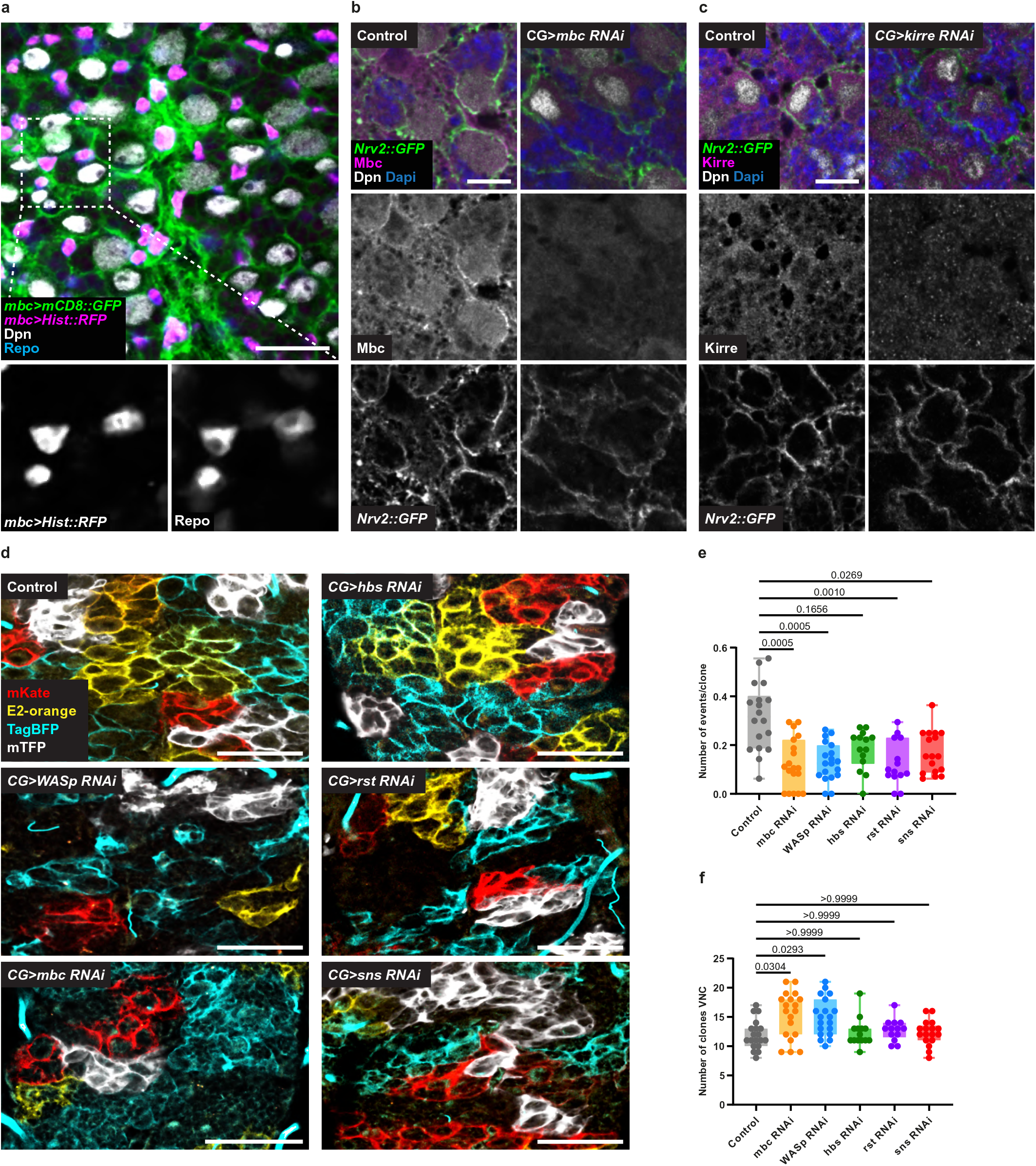
Cell fusion between CG units is regulated by canonical fusion molecules. a) Expression of membrane targeted GFP (*mCD8::GFP*, green) and nuclear RFP (*Hist::RFP*, magenta) using the trojan line *mbc-Gal4* to assess the expression of *mbc* in CG. Glia nuclei were labelled with Repo (blue) and NSC were labelled with Dpn (gray). Scale bar: 20 μm. b) Endogenous expression of Mbc in the CNS assessed by immunostaining with Mbc antibody (magenta) in the VNC. CG membranes are labelled with *Nvr2::GFP* (green), NSC are labelled with Dpn (grey) and Dapi (blue) was used to visualise all nuclei. Upper panels show the expression in control CNS. Lower panels show the expression after RNAi knockdown of *mbc*. Scale bar: 10 μm. c) Endogenous expression of Kirre in the CNS assessed by immunostaining with Kirre antibody (magenta) in the VNC. CG membranes are labelled with Nvr2::GFP (green), NSC are labelled with Dpn (grey) and Dapi (blue) was used to visualise all nuclei. Upper panels show the expression in control CNS. Lower panels show the expression after RNAi mediated down regulation of *kirre*. Scale bar: 10 μm. d) RNAi knockdown of cell-cell fusion related genes in multicoloured labelled CG (Raeppli CAAX) in the VNC. Control (no RNAi), *WASp*, *mbc*, *hbs*, *rst* and *sns* RNAi-knockdowns are shown. RNAi expression was induced at ALH0, larvae were maintained at 29°C and dissected at ALH72. Scale bars: 50 μm. e, f) Quantification of the number of fusion events per clone (e) and number of clones (f) for multicoloured labelled Raeppli CG clones at ALH72 (at 29°C) after knockdown of fusion genes in CG. Results are presented as box and whisker plots. Whiskers mark the minimum and maximum, the box includes the 25th–75th percentile. Individual values are superimposed. Data statistics: one-way ANOVA with a Kruskal–Wallis multiple comparison test.

We then asked whether molecular players associated with myoblast fusion were required for fusion between CG units. We independently knocked down several fusion genes in the CG through RNAi while inducing multicolour clonal labelling (Raeppli-CAAX) and calculated the number of fusion events (overlap between at least two colours, see Methods) per VNC compared to a control condition (Fig. 6e). Strikingly, we observed a significant reduction in the number of fusion events when either *mbc*, *Wasp*, *rst* or *sns* were knocked down (Fig. 6e). For *mbc* and *Wasp*, which showed the most significant reductions, this was paired with a slight increase in number of clones per VNC (Fig. 6f). *hbs*, *kirre* and *lmd* knockdowns also tended towards a reduction in the number of fusion events, albeit the difference was not statistically significant (Fig. 6d-e and Supp. Fig. S6d-f). These data show that known molecular players of classical fusion pathways regulate fusion of CG units.

### Growth and atypical cell-cell fusion are required in CG for correct network architecture and NSC ensheathing

Our results show that CG perform a diversity of cellular processes during niche morphogenesis. Previous studies^72^ had shown that PI3K/Akt-dependent cellular growth was essential to proper network architecture around NSCs (Supp. Fig. S7a, *CG>Δp60*), while preventing mitotic entry through knockdown of *string*/*cdc25* (Supp. Fig. S7a, *CG>stg RNAi*) did not reveal detectable alterations. We enquired about the functional and respective relevance of the different processes we uncovered in building the accurate organisation of the seamless structure of the CG network. As to our knowledge no genetic conditions specifically forcing abscission have been identified in Drosophila so far, we focused on the impact of blocking replication-dependent growth and atypical fusion in CG.

First, we found that knocking down *dup* resulted in dramatic defects in CG growth and network formation (Fig. 7a, *CG>dup RNAi*), with very little CG signal left, which suggested that endoreplication is crucial to CG morphogenesis. The remaining CG cells sometimes harboured a globular morphology, a phenotype reminiscent of blocking membrane vesicular transport in these cells, a condition also associated with loss of proliferation^32^. Accordingly, compared to a control condition, the quantity of CG membrane by NSC was very low, and NSCs were rarely found in individual chambers (Fig. 7b-c).

**Figure 7:**
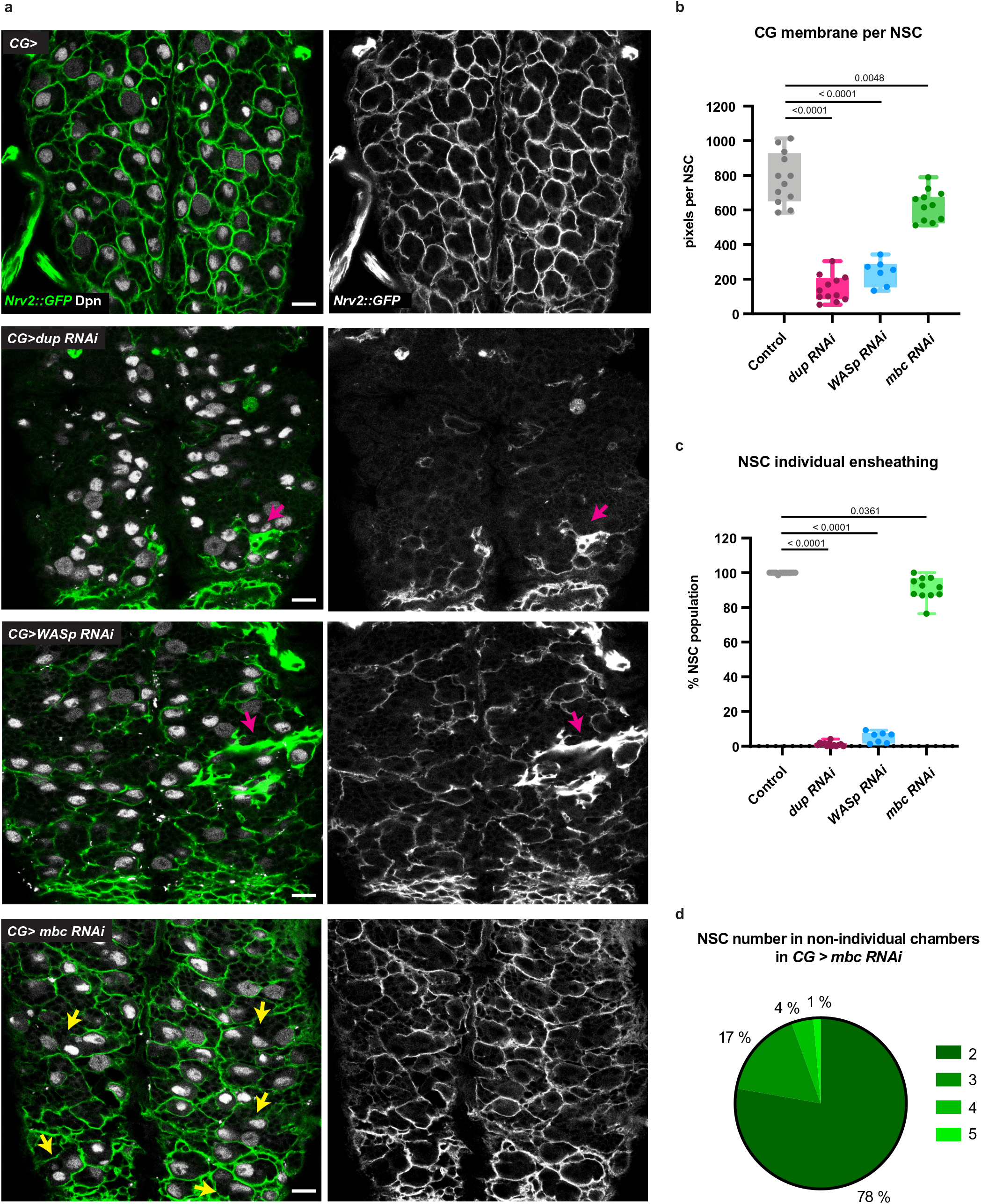
Growth and cell fusion of CG units are required for correct CG architecture. a) Effect of dysregulation of genes involved in endoreplication and atypical fusion on overall CG network architecture. RNAi knockdown of *dup* (DNA replication), *WASp* and *mbc* (cell-cell fusion) are shown (all at ALH72 at 29°C). CG network architecture is visualised with Nrv2::GFP and NSCs are stained with anti-Deadpan (Dpn). Yellow arrows point towards ensheathing of several NSCs (instead of one only) in a chamber of CG membrane. Pink arrows indicate local accumulation of CG membrane. Scale bars: 10 μm. b) Quantification of the average quantity of CG membrane per NSC in the genetic conditions shown in a). See Methods for details. Results are presented as box and whisker plots. Whiskers mark the minimum and maximum, the box includes the 25th–75th percentile. Individual values are superimposed. Data statistics: one-way ANOVA with a Tukey’s multiple comparison test. c) Quantification of the percentage of NSCs individually ensheathed by CG membrane. See Methods for details. Results are presented as box and whisker plots. Whiskers mark the minimum and maximum, the box includes the 25th–75th percentile. Individual values are superimposed. Data statistics: generalized linear model (Binomial regression with a Bernoulli distribution). d) Distribution of the number of NSCs per chamber in non-individual chambers for *mbc RNAi* in CG. Results are presented as a pie chart.

Next, taking advantage of our data identifying molecular regulators of fusion between CG units (Fig. 6), we assessed the impact of their downregulation in the CG. We observed that individually knocking down fusion genes resulted in alterations of the overall CG network structure, ranging in magnitude (Fig. 7a and Supp. Fig. S7b). We first found that *WASp* RNAi led to a striking disorganisation of the CG network, with heterogeneous coverage along the network (Fig. 7a, *CG>WASp RNAi*), less CG membrane available per NSC in average (Fig. 7b) and destruction of NSC chamber structure (Fig. 7c). In addition, we noticed local accumulation of CG membranes (pink arrows, Fig. 7a). Such phenotype was also apparent through Raeppli CAAX (Fig. 6c). As WASp is a general regulator of actin cytoskeleton, by enabling actin nucleation for microfilament branching, it is possible that its effects bypass its strict involvement in fusion mechanisms, leading to strong phenotypes. Looking at other regulators of cell-cell fusion, we observed localised disruptions or alterations in chamber shapes for *mbc*, *sns*, *dock*, and, to a lesser extent, for *kirre* (Supp. Fig. 7b). For *mbc* knockdown in particular, we noticed some heterogeneous distribution of the CG membrane (Fig. 7a, *CG>mbc RNAi*), which was accompanied by a significant decrease in the quantity of CG membrane associated with NSCs (Fig. 7b). Importantly, we also observed several occurrences of CG chambers containing more than one NSC (Fig. 7a, yellow arrows; Fig. 7b), mostly grouped by two (Fig. 7d). This suggests that CG fusion is involved in ensuring the individual ensheathing of NSCs by CG membrane. These observations led us to propose that fusion genes, and especially or at least in a more detectable fashion actin-related genes, are important for the formation of CG network and chamber organisation. All together, these data demonstrate that growth and fusion define the stereotypical architecture of the CG niche both as a network and as a structure of individual ensheathings of NSCs.

## Discussion

Here we dissect the cellular mechanisms supporting the acquisition of architectural complexity in the NSC niche using the morphogenesis of the CG network in Drosophila. We have first uncovered that individual CG cells grow extensively during niche formation. Distinct proliferative strategies convert them into syncytial units in which the different nuclei stay connected, in part through cytoplasmic bridges. We found that these CG units ensheath NSC subsets, covering the entire population in a tile-like fashion. CG units can further undergo homotypic fusion, sharing several subcellular compartments. While this process relies on classical pathways involving conserved cell surface receptors and actin regulators, it is also highly atypical at several levels. Its location is variable, not (yet) predictable, and it is dynamic/transient in time and partial in space, resulting in remodelled compartments from original partners. Ultimately, the combination in time and space of cellular growth, proliferation and fusion are required to build the complex and robust architecture of the CG niche (Fig. 8). Altogether, our findings identify principles of niche formation, revealing unexpected and original cellular processes, while highlighting its impact on organising the NSC population and a remarkable conservation of the spatial partition of glial networks.

**Figure 8:**
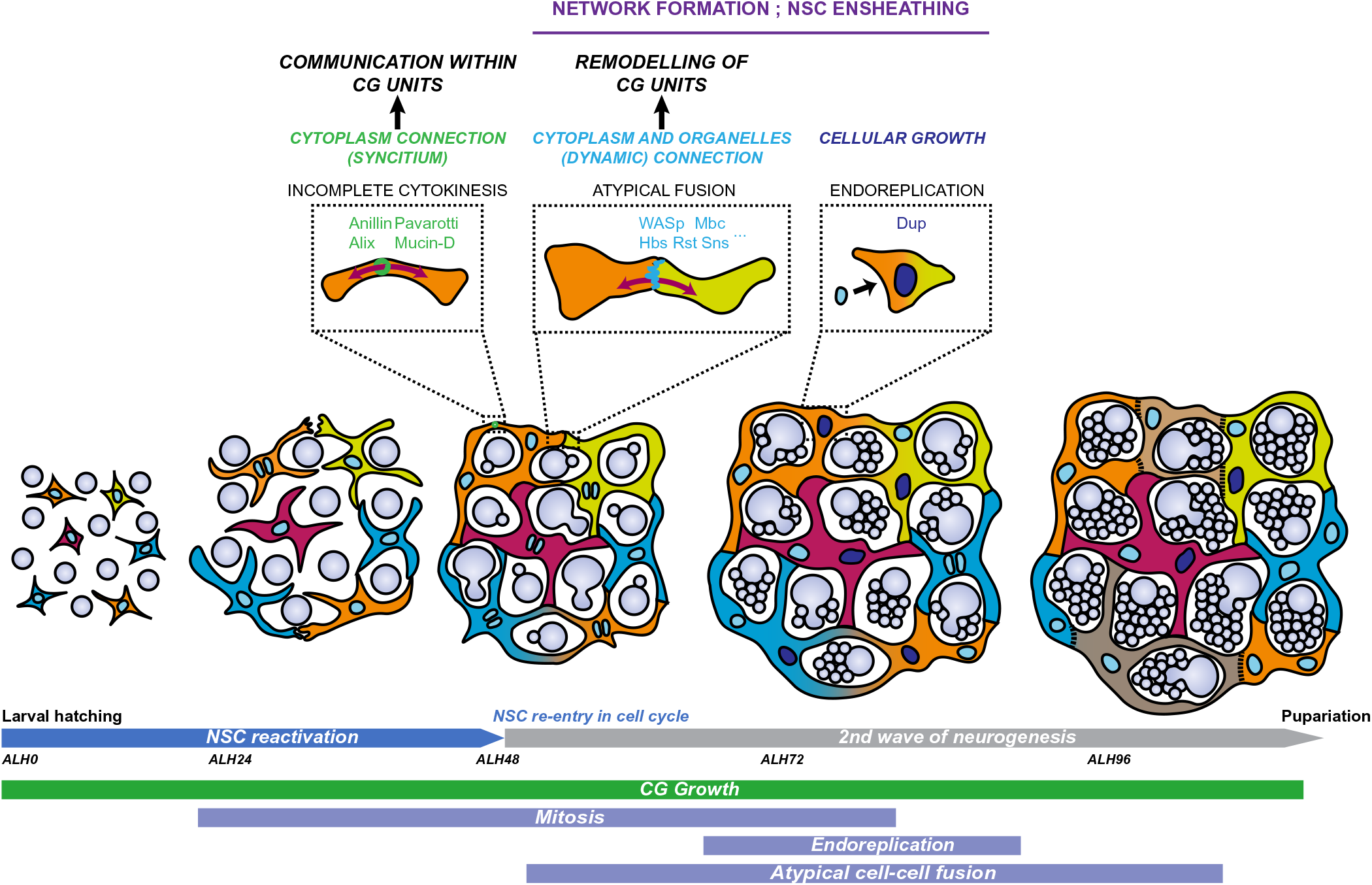
Model of CG morphogenesis along developmental time and NSC behaviour. Individual CG cells grow to tile the CNS, undergoing both endoreplicative and mitotic events that create multinucleated and polyploid cells. These syncytial units are also able to fuse with each other, exchanging subcellular compartments including cytoplasm, membrane and organelles. This fusion appears partial in space and dynamic in time, and can lead to sharp boundaries between connected and unconnected CG domains. Each CG unit is able to enwrap several NSC lineages. Polyploid nuclei are shown in darker blue.

Polyploidy has been associated with large cells or cells that need to be metabolically active, as a way to scale their power of biosynthesis to their cellular functions^39,48^. For example, the megakaryocytes of the bone marrow, which are required to generate large quantities of mRNA and protein for producing platelets, undergo polyploidization. Polyploidy is also an elegant way to support cell growth while protecting a specific cell architecture that would suffer from mitosis-associated adhesion and cytoskeleton changes. In this line, the polyploidization of the subperineurial glia, which exhibit strong junctions to fulfil its role as a blood-brain barrier, maintains barrier integrity in response to CNS growth^45^. The CG cells, which have a highly complex topology integrating NSC position and display large sizes (Fig. 1f) fit both categories.

Importantly, increase in ploidy can be achieved by different processes, many of which rely on variations of the cell cycle^37,38,47,73^, including endocycle, endomitosis and acytokinetic mitosis. Here we propose that CG exhibit several of these cycling strategies. The increase in chromosome number seen in some nuclei (Fig. 2h-i) as well as some aborted DNA segregation at anaphase (Fig. 2l) imply that CG undergo either endocycling or endomitosis without nuclear division. In addition, some CG perform acytokinetic mitosis, displaying all stages of mitosis including midbody formation (Fig. 2c-d, f-g and Supp. Fig. S2b), but without abscission, leading to a syncytial, multinucleated unit of CG cells (Fig. 3c-g). We cannot exclude that some CG cells complete cytokinesis and undergo proper cell division, an outcome challenging to observe considering CG architecture. Interestingly, acytokinetic mitosis takes place in the germline stem cell niche of many animals^74^, including in Drosophila in which the maturing oocyte and supporting nurse cells stay connected by ring canals, intercellular bridges that are stabilized on arrested cleavage furrows^75^. While we identified several components of ring canals in midbody-like puncta present along the CG membrane (Fig. 3e and Supp. Fig. 3b-c), the exact composition and regulation of such structure in the CG remain to be deciphered.

Notably, blocking endoreplication is detrimental to network formation (Fig. 7a-c), whereas preventing the increase in CG nuclei (through knockdown of string/*cdc25*, which prevents mitotic entry, Supp. Fig. S7a or through expression of the cyclin E/cdk2 inhibitor *dacapo*, which blocks G1 to S transition) did not have any detectable impact^27^. This is a puzzling observation suggesting that endoreplication is of higher importance than mitosis in steady-state conditions, and that common players (*i.e*., *dacapo*) might have more instrumental functions in one process versus the other. How the balance between endoreplication and mitosis is regulated, as well as more generally the trigger(s) and timing for these processes are key questions that need to be addressed. The antero-posterior wave of CG cycling (Fig. 2a) is particularly intriguing. Notably, CG proliferation depends on nutrition via activation of the PI3K/Akt pathway^27,40^ (Supp. Fig. S7a). The interplay between spatial and temporal signalling will thus be of special interest.

Using several approaches, including dual and multicolour clonal analysis for different subcellular compartments, FLIP experiments, photoconversion and targeted loss of function, we have shown that CG units have the ability to interact with each other and share their components in a manner dependent of known molecular players of myoblast fusion. A puzzling observation is the spatially-limited nature of this exchange, as witnessed through cytoplasmic and membrane markers (Fig. 4a-c). Our data indeed support the existence of atypical fusion events, partial in space and dynamic in time. Classical cell-cell fusion, such as myoblast fusion, is complete and irreversible, leading to full combination of all components in time. Although some heterogeneity in the mixing of components of the cells of origin could be happening, depending on molecular properties (i.e., membrane proteins; phase separation) or fixed positioning (i.e., nuclei), here we are able to observe sharp boundaries between fused and unfused regions (Fig. 4a-b). A possibility could be that we catch the event at a very early stage. However, in this case we should expect some complete colour overlap at later stages, at least at the same frequency with which partial mixing happened at the previous recorded stages, something we do not see (see Fig. 1e, ALH96, representative of the rarity of complete overlaps at this stage). A fitting explanation could be that the fusion happening between CG cells is somehow transient, and that other, unknown mechanisms exist to rupture and close membranes again, severing the communication between the two original CG units, either on one side or in both. Cytoplasmic exchange between CG units could be constantly remodelling, generating alternating phases of fusion and separation and creating a complex continuum of CG combinations, which could keep evolving over time. Our FLIP experiments (Fig. 5) support this hypothesis by showing that fused domains can lose or alter their connection with the original, still present, CG unit and become a novel cytoplasmic compartment with its own properties. As such, contrary to classical fusion in which two cells lead to one cell/compartment, here two cells can lead to three or more cells/compartments (Fig. 5e). These compartments will inherit characteristics from the original fusing partners, as demonstrated by cytoplasmic mixing and the existence of CG units with connected nuclei of different origins (Fig. 4b-c, e). The observation of a lesser intensity of one of the fluorophores in the shared, fused zone compared to the CG unit of origin (Fig. 4a for membrane and Fig. 5 for cytoplasm) actually fits with the hypothesis of restricted remaining connection with the original CG unit. Interestingly, there has been some previous reports of partial cell fusion (discussed in^76^), suggesting that such phenomenon might be underestimated. The involvement of some of the molecular players controlling myoblast fusion, with conservation in vertebrates (*e.g.*, mbc/Dock180; Kirre/Kirrel) suggests shared adhesion and actin-dependent mechanisms with classical fusion. However, whether similar cell players (*e.g.*, fusion competent cells versus founder cells), molecular interactions and intracellular signalling happen in CG is left to be demonstrated. Recently, full cytoplasmic exchange between cells of the Drosophila rectal papillae have been shown to happen through membrane remodelling and gap junction communication rather than classical fusion pathways^77^.

The parameters regulating the frequency, location and timing of these atypical fusion events also remain mysterious. A way to understand when and where fusion happens might be to understand why it happens. Here we show that fusion between CG units is required for a gapless, seemingly-continuous meshwork as well as for the individual ensheathing of NSCs (Fig. 7a-d). Beyond a more generic role of the actin cytoskeleton in CG architecture, this could suggest that CG fusion somehow ensures that no gap in CG network and in the associated coverage of NSCs is left unmet. Fusion could act as a rescue mechanism, kicking in when seamless tiling between CG units fails. Curiously, CG have a certain capacity to replace each other when ablated, seemingly able to probe space and reach neighbours^32^. How much fusion mechanisms could participate in this sensing and repair is an intriguing question. Another, seducing, hypothesis would see such dynamic fusion as a powerful strategy to modulate the extent of communication and signal exchange within the CG network, as a response either to CG own fluctuating needs or to NSC behaviour, fulfilling its role as a neurogenic niche. The fact that cellular fusion is able to change the number and coverage/size of CG units implies that the spatial, modular partition of the NSC population can be remodelled over time, and possibly upon varying NSC needs. In this line, we noticed slight fluctuations in the number of fusion events (Supp. Fig. S4b), as well as in the number of NSCs encased by one CG unit overtime (Fig. 1h, decrease between ALH72 to ALH96), hinting that remodelling of CG unit boundaries might be a way to control niche properties along neurogenesis. Further work will be needed to assess whether the physical partition of the NSC population also translates into a functional one. This would be crucial to understand how NSCs behave as a coordinated population versus groups of individual cells.

Here we show that a glial network is built from cell growth and fusion mechanisms, resulting in a highly connected, yet partitioned, structure which ensheathes NSCs. These findings uncover principles of niche organisation that ultimately creates a modular structure spatially subdividing the NSC population, a fascinating discovery within the context of individual versus population-based regulation of stem cells. It interesting to note that astrocytes have been shown to set up gap junctions between them, becoming a so-called astrocytic syncytium^78,79^, while at the same time occupying mostly non-overlapping, defined sub-territories^80,81^. Astrocytes in the mammalian NSC niche also form, through their end feet, a reticular structure sitting between neural progenitors and the blood vessels^10^, similarly to the *glia limitans* between the meninges and the cerebral parenchyma^82^. This suggests that connected, modular glial networks might be a common occurrence during CNS development. Understanding the features and regulators of CG morphogenesis, as well as the resulting roles on neurogenesis, thus provides an original blueprint to explore the multi-facetted roles of glial networks, as well as the morphogenetic processes of complex niche structures.

## Methods

### Fly lines and husbandry

Drosophila melanogaster lines were raised on standard cornmeal food at 25°C. Lines used in this study are listed in the table below:

| Strains | Source | Stock number/Reference |
| --- | --- | --- |
| *w<sup>1118</sup>* | BDSC | 5905 |
| *Nervana2::GFP (Nrv2::GFP)* | BDSC | 6828 |
| *tubulin-GAL80<sup>thermosensitive(ts)</sup>* | BDSC | 65406 |
| *Cre recombinase* | BDSC | 851 |
| *yw, hs-FLP* | Andrea Brand lab |  |
| *CoinFLP* | BDSC | 58750 |
| *cyp4g15-GAL4* | BDSC | 39103 |
| *cyp4g15-FRT-STOP-FRT-LexA* | This study |  |
| *cyp4g15-FLP* | This study |  |
| *cyp4g15-mtd::Tomato* | This study |  |
| *alrm-GAL4* | Marc Freeman lab | <sup>83</sup> |
| *mbc-GAL4 (Trojan)* | BDSC | 66840 |
| *UAS-H2B::YFP (Hist::YFP)* | François Schweisguth lab | <sup>84</sup> |
| *UAS-H2B::RFP (Hist::RFP)* | Yohanns Bellaïche lab | <sup>85</sup> |
| *UAS-His3.3.mIFP-T2A-HO1 (Hist::IFP)* | BDSC | 64184 |
| *UAS-GFP* | BDSC | 1522 |
| *UAS-mCD8::GFP* | BDSC | 5130 |
| *UAS-mCD8::RFP* | BDSC | 27399 |
| *UAS-mito::GFP* | BDSC | 8443 |
| *UAS-GFP::Alix* | Jean-René Huynh lab | <sup>54</sup> |
| *UAS-hβactin::ECFP* | BDSC | 7064 |
| *LexAOp-mCherry::mito* | BDSC | 66531 |
| *UAS-Raeppli CAAX 43E* | BDSC, This study | 55082 |
| *UAS-Raeppli NLS 53D* | BDSC, This study | 55087 |
| *LexAop-Raeppli CAAX 43E* | BDSC, This study | 55082 |
| *UAS-mRFP::Scra* | BDSC | 52220 |
| *Fly FUCCI* | BDSC | 55117 |
| *G-TRACE* | BDSC | 28280 |
| *iTRACE* | BDSC | 66387 |
| *Ubi-p63E-GFP::Pavarotti* | David Glover lab | <sup>86</sup> |
| *UAS-mbc RNAi* | BDSC | 32355 |
| *UAS-WASp RNAi* | BDSC | 51802 |
| *UAS-rst RNAi* | VDRC | 27223 |
| *UAS-hbs RNAi* | BDSC | 57003 |
| *UAS-kirre RNAi* | VDRC | 27227 |
| *UAS-Imd RNAi* | BDSC | 42871 |
| *UAS-sns RNAi* | BDSC | 64872 |
| *UAS-dock RNAi* | BDSC | 27728 |
| *UAS-dup RNAi* | BDSC | 29562 |
| *UAS-stg RNAi* | BDSC | 34831 |
| *UAS-Kaede* | BDSC | 26161 |
| *UAS-Δp60* |  | <sup>87</sup> |

### Larval staging

Embryos were collected within 2-4 hours window on grape juice-agar plates and kept at 25°C for 20-24 hours. Freshly hatched larvae were collected within a 1 hour time window (defined as 0 hours after larval hatching, ALH0), transferred to fresh yeast paste on a standard cornmeal food plate and staged to late first instar (ALH24), late second instar (ALH48), mid third instar (ALH72) and late third instar (ALH96).

### DNA cloning and Drosophila transgenics

A portion of the *cyp4g15* enhancer (GMR55B12, Flybase ID FBsf0000165617), which drives in the cortex glia and (some) astrocyte-like glia, was amplified from genomic DNA extracted from *cyp4g15-GAL4* adult flies, with a minimal Drosophila synthetic core promoter [DSCP^88^] fused in C-terminal.

For creating *cyp4g15-FLP*, the *FLP* DNA, which codes for the flippase enzyme, was amplified from the plasmid pMH5^89^ (Addgene 52531). This amplicon together with the *cyp4g15^DSCP^* enhancer were joined using the Multisite gateway system^90^ in the destination vector pDESThaw sv40 (gift from S. Stowers) in order to generate a *cyp4g15^DSCP^-FLP* construct. The construct was integrated in the fly genome at an attP18 docking site through PhiC31 integrase-mediated transgenesis (BestGene). Several independent transgenic lines were generated and tested, and one was kept (*cyp-FLP*).

For creating *cyp4g15-mtd::Tomato*, the *mtd::Tomato* DNA, which codes for a Tomato fluorescent protein tagged at the N-terminal end with Tag:MyrPalm (MGCCFSKT, directing myristoylation and palmitoylation) and at the C-terminal with 3 Tag:HA epitope, was amplified from genomic DNA extracted from QUAS-mtd-Tomato adult flies (BDSC30005, Chris Potter lab), as described in ^91^. This amplicon together with the *cyp4g15^DSCP^* enhancer were joined using the Multisite gateway system^90^ in the destination vector pDESThaw sv40 gift from S. Stowers) in order to generate a *cyp4g15^DSCP^-FLP* construct. The construct was integrated in the fly genome at an attP2 or attP40 docking site through PhiC31 integrase-mediated transgenesis (BestGene). Several independent transgenic lines were generated and tested, and one was kept for each chromosome (*cyp-mtd::Tomato*).

For creating *cyp4g15-FRT-STOP-FRT-LexA*, a FRT STOP cassette was amplified from an UAS-FRT.STOP-Bxb1 plasmid (gift from MK. Mazouni) and the LexA sequence was amplified from the entry vector L2-LexA::p65-L5 (gift from M. Landgraf). The two amplicons were joined together by overlapping PCRs. This *FRT-STOP-FRT-LexA* amplicon together with the *cyp4g15^DSCP^* enhancer were inserted in the destination vector pDESThaw sv40 using Multisite gateway system^90^ to generate a *cyp4g15^DSCP^-FRT-STOP-FRT-LexA::p65* construct. The construct was integrated in the fly genome at an attP2 or attP40 docking sites through PhiC31 integrase-mediated transgenesis (BestGene). Several independent transgenic lines were generated and tested, and one was kept for each docking site.

### Generation of UAS-Raeppli and LexAOp-Raeppli lines

The original construct (BDSC 55082), placing Raeppli CAAX under the control of both UAS and LexAOp sequences, was crossed to a Cre recombinase line (BDSC 851) to randomly excise one of the two control sequences. The resulting lines were checked by PCR to determine whether they carried the UAS or LexAop version.

A similar protocol was followed to generate UAS-Raeppli NLS 53D and LexAOp-Raeppli NLS 53D constructs from the original line BDSC 55087.

### Fixed tissue Immunohistochemistry and imaging

For immunohistochemistry, CNS from staged larvae were dissected in PBS, fixed for 20 min in 4% formaldehyde diluted in PBS with 0.1% Triton X-100, washed two times in PBS-T (PBS+0.3% Triton X-100) and incubated overnight at 4°C with primary antibodies diluted in PBS-T. After washing three times in PBS-T, CNS were incubated overnight at 4°C with secondary antibodies (dilution 1:200) and DAPI (1:1000) diluted in PBS-T. Brains were washed three times in PBS-T and mounted in Mowiol mounting medium on a borosilicate glass side (number 1.5; VWR International). Primary antibodies used were: guinea pig anti-Dpn (1:5000, in-house made, using pET29a-Dpn plasmid from J. Skeath for production), rabbit anti-Dpn (1:200, gift from R. Basto), chicken anti-GFP (1:2000, Abcam ab13970), rat anti-ELAV (1:100, 7E8A10-c, DSHB), mouse anti-Repo 1:100 (DSHB, 8D12-c), rabbit anti-Phospho-histone H3 (1:100, Millipore 06-570), rat anti-mbc^92^ (1/200, gift from S. Abmayr), guinea pig anti-kirre^93^ (1/1000, gift from S. Abmayr) and rabbit anti-Mucin D^94^ (1/1000, gift from AA. Kramerov). Fluorescently-conjugated secondary antibodies Alexa Fluor 405, Alexa Fluor 488, Alexa Fluor 546 and Alexa Fluor 633 (ThermoFisher Scientific) were used at a 1:200 dilution. DAPI (4′,6-diamidino-2-phenylindole, ThermoFisher Scientific 62247) was used to counterstain the nuclei.

### Image acquisition and processing

Confocal images were acquired using a laser scanning confocal microscope (Zeiss LSM 880, Zen software (2012 S4)) with a Plan-Apochromat 40x/1.3 Oil objective. All brains were imaged as z-stacks with each section corresponding to 0.3-0.5 μm. Images were subsequently analysed and processed using Fiji (Schindelin, J. 2012), Volocity 6.3 (Quorum technologies), the Open-Source software Icy v2.1.4.0 (Institut Pasteur and France Bioimaging, license GPLv3) and Photoshop (Adobe Creative Cloud).

### Live imaging

For live imaging, culture chambers were prepared by adding 300 μl of 1% low-melting agarose prepared in Schneider’s medium supplemented with pen-strep on a glass-bottom 35 mm dish (P35G-1.5-14-C, MatTek Corporation) and allowed to solidify. Circular wells of approximately 2 mm diameter were then cut out using a 200 μl pippet tip fitted with a rubber bulb. CNS from staged larvae were dissected in Schneider’s Drosophila medium (21720-024, Gibco) supplemented with 10% heat-inactivated fetal bovine serum (10500, Gibco), penicillin (100 units ml^−1^) and streptomycin (100 μg ml^−1^) (penicillin–streptomycin 15140, Gibco). 4–6 CNS were placed inside small wells of a pre-prepared culturing chamber and covered with culture medium (Schneider’s + 5 % FBS + larval lysate (10 μl/ml) + pen/strep (1/100). Larval lysate is prepared by homogenising twenty 3rd instar larvae in 200 μl of Schneider’s, spinning down once at 6000 rpm for 5min at 4°C, and recovering the supernatant. Brains were set in position and let to settle around 5-10 minutes before starting imaging. Brains were imaged on a laser scanning confocal microscope (Zeiss LSM 880, Zen software (2012 S4)) fitted with a temperature-controlled live imaging chamber (TC incubator for Zeiss Piezo stage, Gataca systems) using a Plan-Apochromat 40x/1.3 Oil objective. Four-dimensional z-stacks of 5–10 μm at 0.5 μm intervals were acquired every 2-3min. Movies were performed on the ventral side of the ventral nerve cord. Images were subsequently analysed and processed using Fiji (Schindelin, J. 2012).

### Quantification of cortex glia nuclei and mitotic cortex glia

Wild-type brains expressing RFP or GFP-tagged (*Hist::RFP* or *Hist::YFP*, respectively) driven by *cyp4g15-GAL4*, were stained with phospho-histone H3 antibody to detect mitotic CG. Entire brains were imaged and quantification of total and mitotic CG nuclei numbers were performed in Volocity 6.3 (Quorum technologies) using adjusted protocols for detection of objects.

### Cell cycle analysis (FUCCI)

We used the Fly-FUCCI system^41^ that allows to discriminate between different phases of the cell cycle by expressing truncated forms of E2F and Cyclin B (CycB) fused to EGFP and mRFP1, respectively (EGFP::E2F 1-230, mRFP1::CycB 1-266). We used the *cyp4g15-GAL4* driver to express UAS-EGFP::E2F 1-230 and UAS-mRFP1::CycB 1-266 in CG cells. Staging of larvae was performed at 25°C and brains were dissected in PBS at ALH0, ALH24, ALH48, ALH72 and ALH96. Brains where immediately fixed in 4 % formaldehide diluted in PBS for 20 min, washed 3 times in PBS and mounted in Mowiol mounting medium on glass slides. Samples were imaged as described above and quantification of G1 (green), S (red) and G2/M CG nuclei was performed in Volocity 6.3 (Quorum technologies).

### Multicolour clonal analyses (Raeppli)

Heat-inducible Raeppli clones were generated by crossing *yw; UAS-Raeppli-CAAX 43E; cyp4g15-Gal4/TM6B* or *yw; UAS-Raeppli-nls 53D; cyp4g15-Gal4/TM6B* males to *hs-FLP* females. For knockdown experiments, chosen RNAi lines were crossed with *yw, hs-FLP; cyp-FRT-STOP-FRT-LexA/CyO; cyp4g15-GAL4, LexO-Raeppli-CAAX 43E*. Freshly hatched larvae (ALH0) were heat shocked for 2 hours at 37°C and aged to ALH24, ALH48, ALH72 and ALH96 at 25°C, or at 29°C for RNAi experiments. For the visualization of clones at ALH0, constitutively expressed *Cyp-FLP* females were crossed to *yw; UAS-Raeppli-CAAX 43E; cyp4g15-Gal4/TM6B* males. Immunolabelling of NSCs for Fig. 1e was performed as described above. For all other experiments, CNS were dissected and fixed for 20 min in 4% formaldehyde in PBS and washed three times in PBS before mounting. Images were acquired as described above using the spectral mode of a Zeiss LSM880 confocal to promote fluorophore separation.

### Quantification of clone volumes (Raeppli)

Raeppli TFP1 clones were chosen for quantification as it is the strongest and sharpest of the four Raeppli fluorophores. Only clones in the ventral nerve cord were measured. Volumes were measured in 3D images using Volocity 6.3 (Quorum technologies).

### Quantification of clone overlap (Raeppli)

Z stacks of Raeppli CAAX 53E clones induced in CG were visualized in Icy v2.1.4.0 (Institut Pasteur and France Bioimaging, license GPLv3). Boundaries of all one-colour clones, for each of the 4 possible, were mapped manually and outlined with polygons. The same was done in the rare case of full colour overlap. Partial overlaps between clones (defined as an overlap between the colours of adjacent clones that do not cover fully any of the two clones) were then counted manually, with their position recorded on the stack by drawing an ellipse. The clones were counted in the VNC only, stopping at the middle of the neuropile coming from the ventral side, as the great majority of NSCs are located ventrally.

The number of overlaps counted corresponds to the number of fusion events, that we then divided by the total number of clones to generate a “Number of events/clones”.

### Clonal analyses using CoinFLP

The recently described Coin-FLP method^62^ was used to generate red and green mosaics of CG cells. CoinFLP clones were generated by crossing *Cyp-FLP; CoinFLP* females to *yw; LexAop-mCherry; UAS-GFP* or *yw; LexAop-mCherry::mito; UAS-mito-GFP* males and maintained at 25°C. Larvae were staged to ALH48-ALH72 at 25°C. For fixed tissue analyses, brains were dissected and fixed for 20 min in 4% formaldehyde in PBS and washed three times in PBS before mounting. Images were acquired as described above. For live imaging and FLIP experiments (see below), CNS were dissected in Schneider’s medium and mounted as described for live imaging.

### Fluorescence loss in photobleaching (FLIP)

FLIP experiments were performed in dissected larval brains mounted as described above for live imaging. Fluorescence in a selected region of interest (ROI) within a CG cell was repeatedly photobleached over time, and loss of fluorescence in nonbleached regions were monitored. Bleaching was performed on GFP expressed in CG using the *cyp4g15-GAL4* driver. Laser line 488 was used at 100%. Images were acquired as follows: one z-stack of 5–10 μm at 0.3-0.5 μm intervals before bleaching (Pre-bleach), followed by 100 continuous acquisitions at the bleaching plane during the bleaching (Bleach) and one z-stack of 5– 10 μm at 0.3-0.5 μm intervals after bleaching (Post-bleach). Images were subsequently analysed and processed using Fiji.

### Quantification of Fluorescence loss in photobleaching (FLIP)

Measures of fluorescence intensities over time (Fig. 5 and Supp. Fig. 5-1c) were performed on Volocity 6.3 (Quorum technologies). For each region (GFP only, mCherry only, fused H_GFP_ and fused L_GFP_), a ROI was drawn manually to follow the contours of the corresponding area at time T0. The same ROI was kept throughout, except for Fig. 5d, in which a slight x-shift of the ROI shape was performed at the last timepoint (T100) to adjust a restricted x-drift in the tissue. Mean intensities (I_MEAN_) were calculated for each channel in each ROI. The percentage of fluorescence loss in the ROI for each channel (%FL) was determined with the following formula: %FL = (I_MEAN_ Tstart – I_MEAN_ Tend) / I_MEAN_ Tstart.

Due to the existence of i) FLIP-independent decay in fluorescence due to imaging-related photobleaching, and ii) of FLIP-independent variations in fluorescence over time (*e.g.*, small tissue z-shifts, intracellular movements) as well as iii) potential unknown effects of photobleaching of one fluorophore on the other one, we determined for each fluorophore the maximum percentage of loss in fluorescence under which it can be attributed to random variations, with a confidence level of 95%. %FL during a FLIP experiment exceeding this maximum can therefore be considered as a significant variation.

To do so, for each movie, we performed a Monte-Carlo analysis on %FL in the channel corresponding to the unbleached fluorophore (Figures 5a: mCherry; 5b: GFP; 5c: GFP and 5d: mCherry; python script available upon request). This was achieved by sampling %FL over ten thousand randomly positioned 10 × 10 μm squares. A Cumulative Distribution Function was then generated from the results and used to calculate the %FL value required for a 99% confidence level (*i.e.*, if %FL exceeds this value, it is unlikely to be due to random variations, but can be attributed to the FLIP photomanipulation). For each fluorophore, as we treated two movies, we obtained two %FL values fitting the 95% confidence level (Supp. Fig. 5-1b). We then kept the most stringent (*i.e.*, higher) one. For GFP, the threshold is 19.1% and for mCherry, the threshold is 20.8%. As such, only %FL values above these thresholds were considered significant during the FLIP experiments for attributing the loss in fluorescence to the FLIP.

### Kaede photoconversion

Photoconversion experiments were performed in dissected larval brains mounted as described above for live imaging. GFP fluorescence in a selected region of interest (ROI) within a CG cell was illuminated with iterative pulses (each cycle) of a 405 nm diode. Diode power was between 3 and 4%. While single pulse achieved localised conversion in the ROI, it was not enough to visualize diffusion of the converted Kaede form in the CG units, which are of large size.

Images were acquired as follows: one z-stack of 30-40 μm at 0.5-1 μm intervals before photoconversion (Pre-photoconversion), followed by 50 continuous acquisitions at the bleaching plane during the photoconversion (Photoconversion) and one z-stack of 30-40 μm at 1 μm intervals after photoconversion (Post-photoconversion). Images were subsequently analysed and processed using Fiji.

For visualizing Raeppli-NLS and Kaede simultaneously, we used spectral imaging (Zeiss Quasar 34 channels) to acquire and distinguish between mTFP1, GFP, mOrange, mRFP and mKate.

### Quantitative analysis of ploidy by fluorescence in situ hybridization (FISH) of chromosomes

The FISH protocol was performed as previously described^44^ using oligonucleotide probes for chromosomes II and III labelled with 5′CY3 and FAM488 fluorescent dyes respectively (gift from R. Basto). FISH was performed in CNS expressing Histone::RFP or Histone::GFP in CG and dissected in PBS at ALH0, ALH24, ALH48, ALH72 and ALH96. Briefly, dissected brains were fixed for 30 min in 4% formaldehyde prepared in PBS with 0.1% tween 20, washed three times/ 10min in PBS, washed once 10min in 2xSSCT (2xSSC (Sigma S6639) + 0.1% tween-20) and once in 2xSSCT 50% formamide (Sigma 47671). For the pre-hybridization step, CNS were transfered to a new tube containing 2xSSCT 50% formamide pre-warmed at 92°C and denatured 3min at 92°C. For the hybridization step, the DNA probe (40-80 ng) was prepared in hybridization buffer (20% dextran sulphate, 2XSSCT, 50% deionized formamide (Sigma F9037), 0.5 mg ml−1 salmon sperm DNA) and denatured 3min at 92°C. Probes were added to the brains samples and hybridize 5min at 92°C followed by overnight hybridization at 37°C. Samples were washed with 60°C pre-warmed 2XSSCT for 10 min, washed once 5min in 2XSSCT at RT and rinsed in PBS. CNS were mounted in Mowiol mounting medium and imaged as described above. FISH signals for chromosomes II and III were quantified in randomly selected CG nuclei using adapted protocols for dots inside objects detection in 3D images in Volocity 6.3 (Quorum technologies).

### Cortex glial membrane measurements

Each VNC was sampled with six cubes (x = 150 pixels; y = 150 pixels; z = until the neuropile) devoid of trachea or nerve signal. NSC numbers within each cube were determined manually, and the CG membrane signal (using Nrv2::GFP as a proxy) was segmented using a HK-means thresholding (Icy v2.1.4.0,with k = 2). The sum of selected pixels divided by NSC number defines the ratio of CG membrane to NSCs for each cube. A mean from the six cubes was calculated for each VNC, giving an estimation of the ratio of CG membrane per NSC within one brain. The different conditions were analysed via a one-way ANOVA.

### Quantification of individual NSC ensheathing

For each VNC the total number of NSCs was determined through HK-means segmentation in the corresponding channel (Icy v2.1.4.0) and corrected manually. When most of the NSCs did not appear individually encased (*dup RNAi*; *WASp RNAi*), the remaining number of NSCs that were still individually ensheathed by CG membrane was counted manually. For conditions in which most NSCs were still individually ensheathed (control; *mbc RNAi*), we recorded the number of chambers with more than one NSC, as well as the number of NSCs within each. This allowed us to subtract the number of NSCs not individually encased from the total NSC population. Ultimately, the ratio between the number of NSCs individually ensheathed and the total NSC population defines the percentage of individual NSC ensheathing. Taking in consideration the non-normal distribution of the control, the significance of each condition compared to control was then determined through a generalized linear model (Binomial regression with a Bernoulli distribution).

### Statistics and reproducibility

Statistical tests used for each experiment are stated in the figure legends. Statistical tests were performed using GraphPad Prism 7.0a.

## Acknowledgments

We thank the Abmayr and Workman labs for the generous gift of the anti-Mbc and anti-Kirre antibodies; the Basto lab for the DNA FISH probes and the rabbit anti-Dpn; AA Kramerov for the gift of the anti-Mucin-D antibody; Steven Stowers and Matthias Landgraf for sharing plasmids for Multisite Gateway cloning; M’hamed Khalil Mazouni for the sequence of the FRT-STOP-FRT cassette; the Freeman lab for *alrm-GAL4*; James Skeath for the Dpn plasmid; the Bloomington Drosophila Stock Center and the Vienna Drosophila Research Center for RNAi lines. We are grateful to Mateusz Trylinski for help with Fiji scripts and discussions for ploidy count; to Jean-Yves Tinevez (Image Analysis Hub, Institut Pasteur) and Francis Murphy for help with image processing and FLIP quantification; and to Elise Jacquemet (Biostatistics Hub, Institut Pasteur) for help with statistical analysis. Julie Marc generated part of Fig. 1E. Laurence Arbogast built the *cyp4g15-FLP*, *cyp4g15-mtd::Tomato* and *cyp4g15-FRT-STOP-FRT-LexA* constructs. We thank Jean-René Huynh, Juliette Mathieu and Romain Levayer for critical reading of the manuscript.

This work has been funded by a starting package from Institut Pasteur/LabEx Revive, a JCJC grant from Agence Nationale de la Recherche (NeuraSteNic, ANR-17-CE13-0010-01) and a Projet Fondation ARC from Fondation ARC pour la Recherche sur le Cancer to PS. MAR has been supported by a LabEx Revive postdoctoral fellowship and BD by an Amgen fellowship.

## Author Contributions

MAR performed all experiments, except for: Figure 3f-g, Supp. Figure S3d, Figure 4e, Supp. Figure S4f, Figure 6a-c and Supp. Figure S6b-c performed by DB; Figure 3c-d, and Supp. Figure S1d-e performed by BD under the supervision of MAR; and Figures 3e, Fig. 7, Supp. Figure S3b-c, Supp. Figure S4d-e, and Supp. Figure S7a-b performed by PS. MAR, BD and PS quantified and analysed the data. MAR and PS wrote the article and made the figures.

## Declaration of Interests

The authors declare no competing interest.

## Data availability statement

The datasets generated during and/or analysed during the current study are available from the corresponding author on reasonable request.

## Notes

### Competing Interest Statement

The authors have declared no competing interest.

