## Supplemental figures and legends for "An interplay between cellular growth and atypical fusion defines morphogenesis of a modular glial niche"

**Supplemental legends Rujano et al.**

**Supp. Fig. S1**

a) Upper panels show a side view of the CNS showing the astrocyte glia (*alrm>CD8::RFP*, magenta) compartment compared to Nrv2::GFP (green) expression (upper panels). Lower panels are side views of a CNS expressing membrane targeted CD8 (*mCD8::GFP*, green) and Histone (*Hist::RFP*, magenta) in the CG (*cyp4g15-GAL4*). Note that CG are ventral in the VNC, paralleling NSC location, while the astrocyte glia are restricted to the dorsal side. Scale bars: 50 µm.

b) Schematics of the Raeppli multicolour lineage tracing tool. Adapted from [1]. The Raeppli construct contains 5X UAS and LexO sites that act on a basal *hsp70* promoter (cyan ellipse) for expression of chosen fluorescent proteins. UAS and LexO sites are flanked by Lox2272 or LoxP sites, respectively. Cre protein excises one of the enhancers in a mutually exclusive manner. A single attB site (light grey arrow) downstream of the enhancer and promoter, is available for recombination with one of the attP sites (dark grey arrows). A full *hsp70* promoter (dark blue ellipse) regulates the expression of the integrase gene which is blocked by the presence of a stop cassette (black box) flanked by FRT sites (purple triangles). Fluorescent protein genes have a stop codon and are arranged in a linear fashion each preceded by an attP site. Expression of the flippase (Flp) removes the stop cassette and triggers expression of integrase. Heat shock then causes the expression of the integrase gene and the Integrase protein recombines the attB site with one of the attP sites of the construct and removes the in between region. The remaining fluorescent protein is then expressed by the action of LexA or Gal4. Since each fluorescent protein gene is followed by a stop codon, the only expressed fluorescent protein is the one brought closest to the promoter.

c) Progressive increase in CG nuclei numbers visualized at ALH0, ALH24, ALH48, ALH72 and ALH96 (at 25°C). CG nuclei are labelled with *Hist::RFP* driven by *cyp4g15-GAL4*. Scale bars: 50 µm.

d) Quantification of CG nuclei number in the CNS at ALH0 (n=8), ALH24 (n=11), ALH48 (n=8), ALH72 (n=4) and ALH96 (n=7). n, number of CNS. Results are presented as box and whisker plots. Whiskers mark the minimum and maximum, the box includes the 25th–75th percentile, and the line in the box is the median. Individual values are superimposed. Data statistics: ordinary one-way ANOVA with a Tukey's multiple comparison test.

e, f) Quantification of CG nuclei in Raeppli clones induced in CG at ALH0 (e) and assessed at ALH24 (n=9), ALH48 (n=76), ALH72 (n=46) and ALH96 (n=42) at 25°C. n, number of clones. The pan-glial marker Repo (magenta) was used to identify the CG nuclei within clones. Scale bars: 50 mm. The quantification (f) was performed in the brighter clones (mTFP1, cyan) to facilitate earlier time points assessment. The image on the left shows close-up of such clones and their contained nuclei. Results are presented as box and whisker plots. Whiskers mark the minimum and maximum, the box includes the 25th–75th percentile, and the line in the box is the median. Individual values are superimposed. Data statistics: ordinary one-way ANOVA with a Tukey's multiple comparison test.

**Supp. Fig. S2**

a) Schematics of the Drosophila Fucci tool, adapted from [2]. The colours used in our scheme, reflect the colours used in Figures 2a,b and 3b. In early M phase, both GFP-E2F1$_{1–230}$ (green) and mRFP1-CycB$_{1–266}$ (magenta) are present thus labelling the cells grey. Midway mitotic stage, the APC/C marks mRFP1-CycB$_{1–266}$ for proteasomal degradation leaving the cells fluorescing green due to GFP-E2F1$_{1–230}$ expression. As cells progress from G1 to S phase, CRL4$^{Cdt2}$ degrades GFP-E2F1$_{1–230}$, and cells are thus labelled in magenta, because only mRFP1-CycB$_{1–266}$ is present. After cells enter G2 phase, GFP-E2F1$_{1–230}$ protein levels reaccumulate, marking the cells grey due to the presence of mRFP1-CycB$_{1–266}$.

b) Still images of a time-lapse movie (Movie S2) of mitotic CG expressing Hist::RFP (magenta) to label nuclei and Lamin::GFP (green) to label the nuclear envelope. Scale bar: 5 µm.

c) Expression of mRFP::scra (magenta) in CG to monitor midbodies along CG membranes. CG membranes and nuclei are labelled with Nrv2::GFP (green) and Hist::IFP (blue) respectively. Arrows indicate midbodies. Scale bar: 10 µm.

**Supp. Fig. S3**

a) Sharing of cytoplasmic material between CG assessed by Fluorescence Loss In Photobleaching (FLIP) of cytosolic GFP (green). Top panels depict a region in the VNC before (pre-bleach) and after bleaching (post-bleach). CG nuclei are labelled with Hist::RFP (magenta). Bottom panels show intermediate time points (GFP only, pseudocolored with thermal LUT) during continuous photobleaching. Bleached area is delineated by the dashed square. Scale bars: 20 µm.

b) Puncta containing both anillin (mRFP::scra, magenta ) and Mucin-D (anti-Mucin-D, cyan), are localising along the CG membrane (Nrv2::GFP, green). CG nuclei are stained with His::IFP (*CG>His::IFP*, grey). Scale bars: 2 µm.

c) Puncta enriched both in Mucin-D (anti-Mucin-D, grey) and Pavarotti (Ubi-p63E-GFP::Pavarotti, green), two classical components of midbodies and intercellular bridges, are found along the CG membrane (*cyp4g15-mtd::Tomato*, magenta). Scale bars: 2 μm.

d) CG connection via the midbodies marked by anillin (mRFP::scra, magenta) assessed by FLIP of cytosolic GFP (green). Top panels depict a region in the VNC before (pre-bleach) and after bleaching (post-bleach). CG nuclei are labelled with *Hist::RFP* (magenta). The bleached area delineated by the white dashed square is placed close to an isolated midbody (clear blue inset) in between CG cells. Bottom panels show intermediate time points (GFP only, pseudocolored with thermal LUT) during continuous photobleaching. Scale bars: 10 μm.

**Supp. Fig. S4**

a) Quantification of the number of Raeppli clones at ALH24 (n=5), ALH48 (n=12), ALH72 (n=9) and ALH96 (n=9) at 25°C. n, number of CNS analysed. Results are presented as box and whisker plots. Whiskers mark the minimum and maximum, the box includes the 25th–75th percentile, and the line in the box is the median. Individual values are superimposed. Data statistics: ordinary one-way ANOVA with a Tukey's multiple comparison test.

b) Quantification of colour overlap events in Raeppli clones at ALH24 (n=5), ALH48 (n=12), ALH72 (n=9) and ALH96 (n=9) at 25°C. n, number of CNS analysed. Results are presented as box and whisker plots. Whiskers mark the minimum and maximum, the box includes the 25th–75th percentile, and the line in the box is the median. Individual values are superimposed. Data statistics: ordinary one-way ANOVA with a Tukey's multiple comparison test.

c) Schematics of the Drosophila Coin-FLP technique adapted from [3]. *CoinFLP-Gal4* used with a *FLP*-expressing enhancer triggers the recombination between either the canonical *FRT* sites, resulting in excision of the STOP cassette and expression of LexGAD, or between the *FRT3* sites, that results in excision of the stop cassette and Lex-GAD, thus triggering Gal4 expression.

d) Visualisation of neighbouring Raeppli clones using the nuclear targeted Raeppli constructs at ALH24, ALH48, ALH72 and ALH96 at 25°C. Scale bars: 50 mm.

e) Higher magnification of neighbouring nuclear Raeppli clones that share nuclear material. ALH48 at 25°C. Scale bar: 5 μm.

f) Continuity between CG units due to cellular fusion was assessed by photoconversion of cytosolic Kaede expressed in the CG in combination with early induction of multicolour labelling of CG nuclei (Raeppli-NLS) that leads to clonal labelling of the nuclei in CG units.

Iterative photoconversion was performed in a small area (dashed rectangle) within a Raeppli-NLS CG clone containing nuclei of one colour. Top panels depict the assessed area before (pre-photoconversion) and after photoconversion (post-photoconversion). Bottom panels show the converted form (cKaede) only, pseudocolored with thermal LUT before and after photoconversion, with nuclei represented by black discs outlined in the respective Raeppli colour. In total, two different colours of nuclei are joined by the cKaede signal. Scale bars: 10 $\mu$m.

g) Continuity between CG units due to cellular fusion was assessed by FLIP of CG expressing cytosolic GFP in combination with early induction of multicolour labelling of CG nuclei (Raeppli-NLS) that leads to clonal labelling of the nuclei in CG units. Continuous bleaching was performed in a small area (dashed rectangle) containing nuclei of one colour. Top panels depict the assessed area before (pre-bleach) and after bleaching (post-bleach). Bottom panels show intermediate time points (GFP only, pseudocolored with thermal LUT) during continuous photobleaching. Scale bars: 20 µm.

**Supp. Fig. S5, Supplement 1**

a) Different zones defined by different combinations of GFP and mCherry levels as well as by positions are identified in the area imaged in Figure 5a-d. Left panel, borders of the zones drawn on a still image of experiment 5a (Movie S7) at T0. Right panel, schematic of the area overlaid with the borders and names of the different zones.

b) Estimation of the values that %FL should take to be significantly due to the FLIP experiment rather than chance. For each movie, we performed a Monte-Carlo analysis on %FL in the channel corresponding to the unbleached fluorophore (Figures 5a: mCherry; 5b: GFP; 5c: GFP and 5d: mCherry). This was achieved by sampling %FL (percentage of change in mean intensity) over ten thousand randomly positioned 10 X 10 $\mu$m squares. Histograms (left panels) represent the distribution of %FL amongst the 10.000 random squares. Cumulative Distribution Functions (middle panels) were generated from the results and used to calculate the %FL value required for a 95% confidence level (right panels).

c) Table of the changes in mean intensities ($I_{MEAN}$) in GFP and mCherry in the different zones of interest between T0 (start of the movie) and T100 (end of the movie/photobleaching), and the corresponding %FL values. The significant %FL values take the colour (green or magenta) of the bleached fluorophores, and the ones outside of the zone targeted by the bleach are in a pale yellow background. See Methods for details on the significance.

**Supp. Fig. S5, Supplement 2**

a-d) Individual GFP and mCherry channels before (pre-bleach) and after bleaching (post-bleach), corresponding to experiments shown in Figure 5a-d respectively. Dashed lines indicate the zones outside of the region that was targeted by the FLIP in which a significant %FL occurred. Scale bars: 10 μm.

e) Still images of time-lapse movies from three different VNCs (ALH70 at 25°C) expressing a fusion of human β-actin with ECFP in the CG only (*cyp>β-actin::ECFP*, grey). Yellow arrows indicate area of high actin remodelling. Scale bars: 10 μm.

**Supp. Fig. S6**

a) Schematics of cell-cell fusion based on the model of myofibers formation in Drosophila, adapted from [4–6]. In this model, a fusion-competent cell and a founder cell recognize and bind to each other, creating a so-called fusogenic synapse. Key, well-characterized players for this step are the cell recognition and adhesion molecules that mediate the binding between the two membranes. These molecules are differentially localised in the fusing cells, expressed either by the fusion competent cell (Sns and Hbs) or by the founder cell (Kirre/Duf and Rst). Binding between partners initiate intracellular signalling, through adapter proteins that will lead to remodelling of the actin cytoskeleton in both cells. In the fusion competent cell, the combined actions of multiple actin regulators (WASp, Rac, Scar, Arp2/3) generate invasive podosome-like protrusions at the interface with the founder cell. These structures trigger a Myosin II- and spectrin-mediated response in the founder cell, which is followed by hemifusion of membranes, pore formation and expansion, culminating in the creation of a multinucleated cell.

b, c) Expression of the lineage tracing tools i-TRACE (b) and G-TRACE (c) under *mbc-GAL4* (Trojan line) to assess *mbc* expression in the CG throughout development. Images were taken at ALH72. Scale bars: 50 μm. Lower panels are higher magnifications of the regions in the dashed squares. Scale bars: 10 μm.

d) RNAi knockdown of cell-cell fusion related genes *kirre* and *lmd* in multicoloured labelled CG in the VNC. RNAi expression was induced at ALH0, larvae were maintained at 29°C and dissected at ALH72. Scale bars: 50 μm.

e, f) Quantification of the number of fusion events per clone (e) and number of clones (f) in multicoloured labelled Raeppli CG clones at ALH72 (at 29°C) after *kirre* and *lmd* knockdown in CG. Results are presented as box and whisker plots. Whiskers mark the minimum and maximum, the box includes the 25th–75th percentile. Individual values are superimposed. Data statistics: one-way ANOVA with a Kruskal–Wallis multiple comparison test.

**Supp. Fig. S7**

a) Effect of overexpression of Δp60 (blocking PI3K/Akt-dependent growth) and of RNAi knockdown of *stg* (blocking entry in the cell cyle) on CG network architecture (visualised with Nrv2::GFP, portion of the VNC). All ALH72 at 29°C. Scale bar: 10 µm.

b) Effect of down regulation of cell-cell fusion gene *mbc*, *sns*, *kirre*, *dock* and *lmd* on CG network architecture (portion of the VNC). CG network architecture is visualised with Nrv2::GFP. All ALH72 at 29°C. Blue arrows point towards localised defects in the network architecture. Pink arrows indicate local accumulation of CG membrane. Scale bar: 10 µm.

**Supp. Movie S1**

Time-lapse movie of mitotic CG expressing *Hist::RFP* (grey).

**Supp. Movie S2**

Time-lapse movie of mitotic CG expressing Hist::RFP (magenta) to label nuclei and Lamin::GFP (green) to label the nuclear envelope. Individual channels are shown in grayscale.

**Supp. Movie S3**

Time-lapse movie of a CG expressing *Hist::RFP* (grey) undergoing endomitosis.

**Supp. Movie S4**

Time-lapse movie of two CG expressing *Hist::RFP* (grey) undergoing mitosis synchronously.

**Supp. Movie S5**

Combined z-stacks before (left) and after (right) photoconversion of a region in the VNC in which CG express the photoconvertible cytoplasmic Kaede (GFP, green, to RFP, magenta) as well as fluorescently marked anillin (mRFP::scra, magenta). mRFP::scra also localises in the nuclei. The cKaede signal overlaps with several midbody-like structure (mRFP::scra puncta, dashed white circles) throughout the z-stack, showing that CG units are rich in intercellular bridges. Scale bar: 10 µm. z-step is 0.50 µm. ALH72, 25°C.

**Supp. Movie S6**

Combined z-stacks before (left) and after (right) photoconversion of a region in the VNC in which CG express the photoconvertible cytoplasmic Kaede (GFP, green, to RFP, magenta) in combination with early induction of multicolour labelling of CG nuclei (Raeppli-NLS, blue, grey and orange) that leads to clonal labelling of the nuclei in CG units. Scale bars: 10 μm. z-step is 1 μm. ALH72, 25°C.

**Supp. Movie S7**

Propagation of information/signals between fused areas was assessed by FLIP in clones generated by CoinFLP with cytosolic GFP (green) and mCherry (magenta) in CG. A GFP expressing clone with areas of partial and no overlap with mCherry was selected. Time-lapse of continuous bleaching of GFP in the non-overlapping part of the GFP clone. Scale bar: 5 μm. ALH48, 25°C.

**Supp. Movie S8**

Propagation of information/signals between fused areas was assessed by FLIP in clones generated by CoinFLP with cytosolic GFP (green) and mCherry (magenta) in CG. A GFP expressing clone with areas of partial and no overlap with mCherry was selected. Time-lapse of continuous bleaching of mCherry in the overlapping part of the GFP clone with high GFP intensities ($H_{GFP}$ subzone). Scale bar: 5 μm. ALH48, 25°C.

**Supp. Movie S9**

Propagation of information/signals between fused areas was assessed by FLIP in clones generated by CoinFLP with cytosolic GFP (green) and mCherry (magenta) in CG. A GFP expressing clone with areas of partial and no overlap with mCherry was selected. Time-lapse of continuous bleaching of mCherry in the overlapping part of the GFP clone with lower GFP intensities ($L_{GFP}$ subzone). Scale bar: 5 μm. ALH48, 25°C.

**Supp. Movie S10**

Propagation of information/signals between fused areas was assessed by FLIP in clones generated by CoinFLP with cytosolic GFP (green) and mCherry (magenta) in CG. A GFP expressing clone with areas of partial and no overlap with mCherry was selected. Time-lapse of continuous bleaching of GFP overlapping part of the GFP clone with lower GFP intensities ($L_{GFP}$ subzone). Scale bar: 5 μm. ALH48, 25°C.

**Supp. Movie S11**

Time-lapse movie of the region of interaction between two neighbouring CG clones generated with CoinFLP and expressing either cytosolic GFP or mCherry, at ALH48.

Rujano et al. Figure 1. Supplement 1

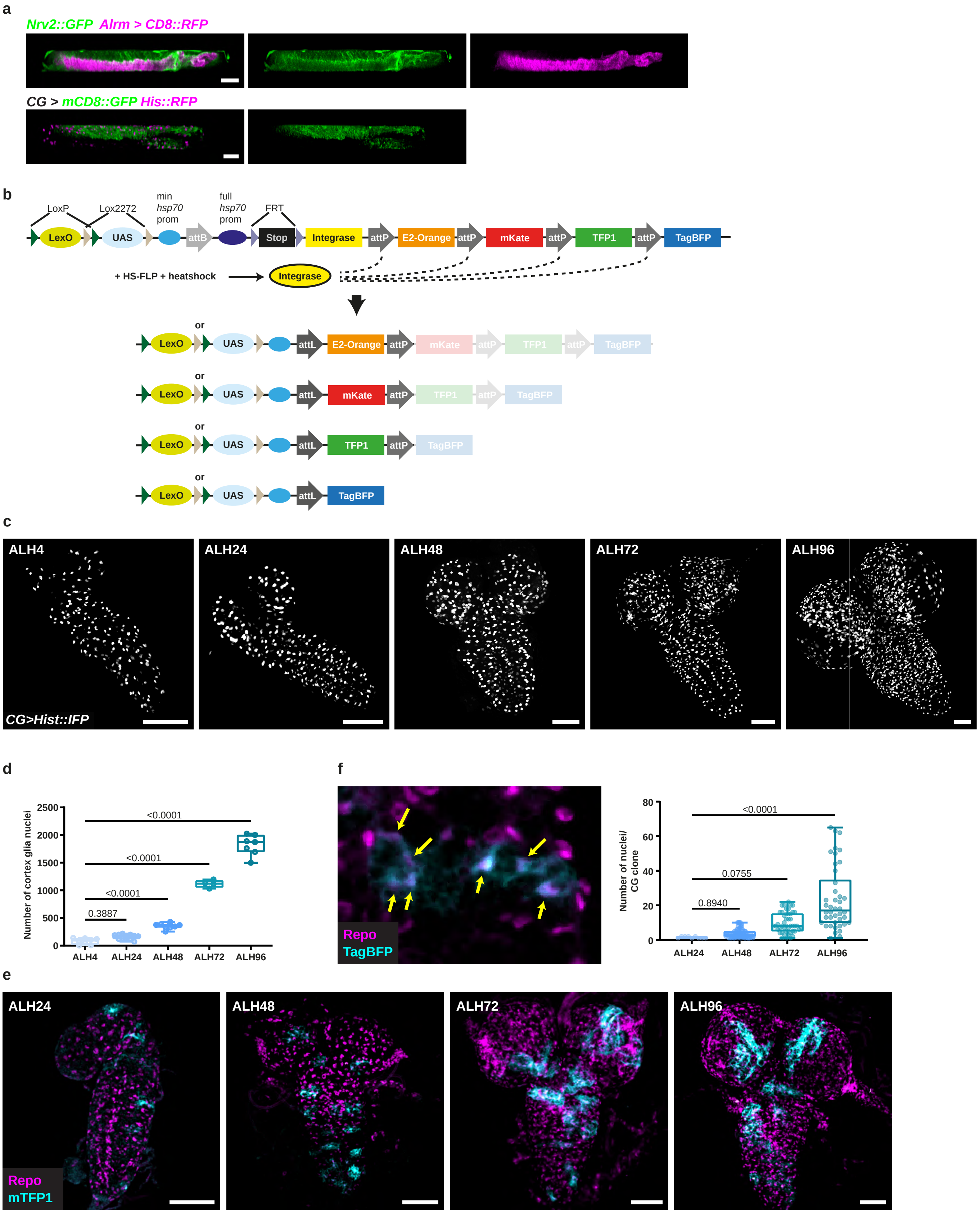

Rujano et al. Figure 2. Supplement 1

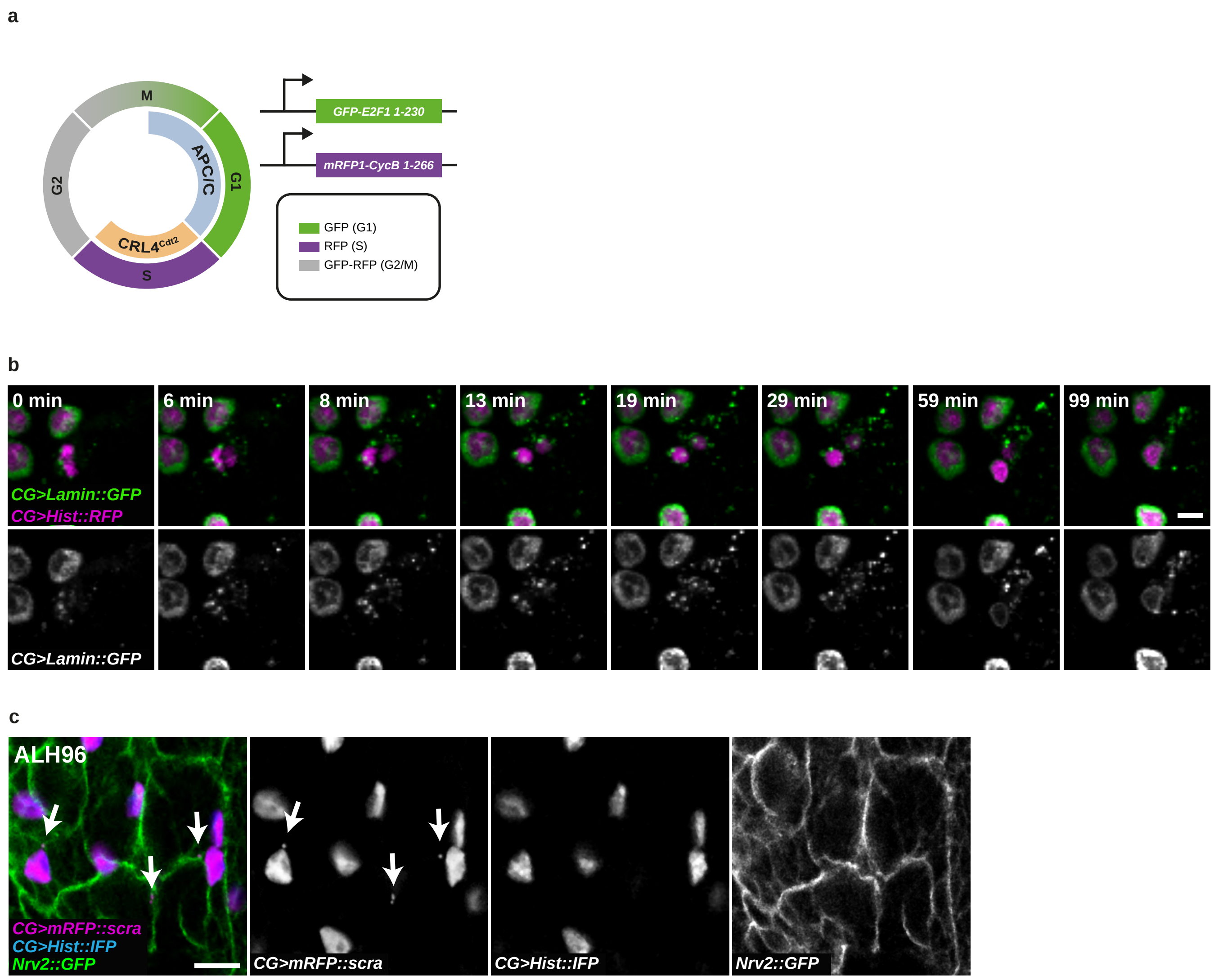

Rujano et al. Figure 3. Supplement 1

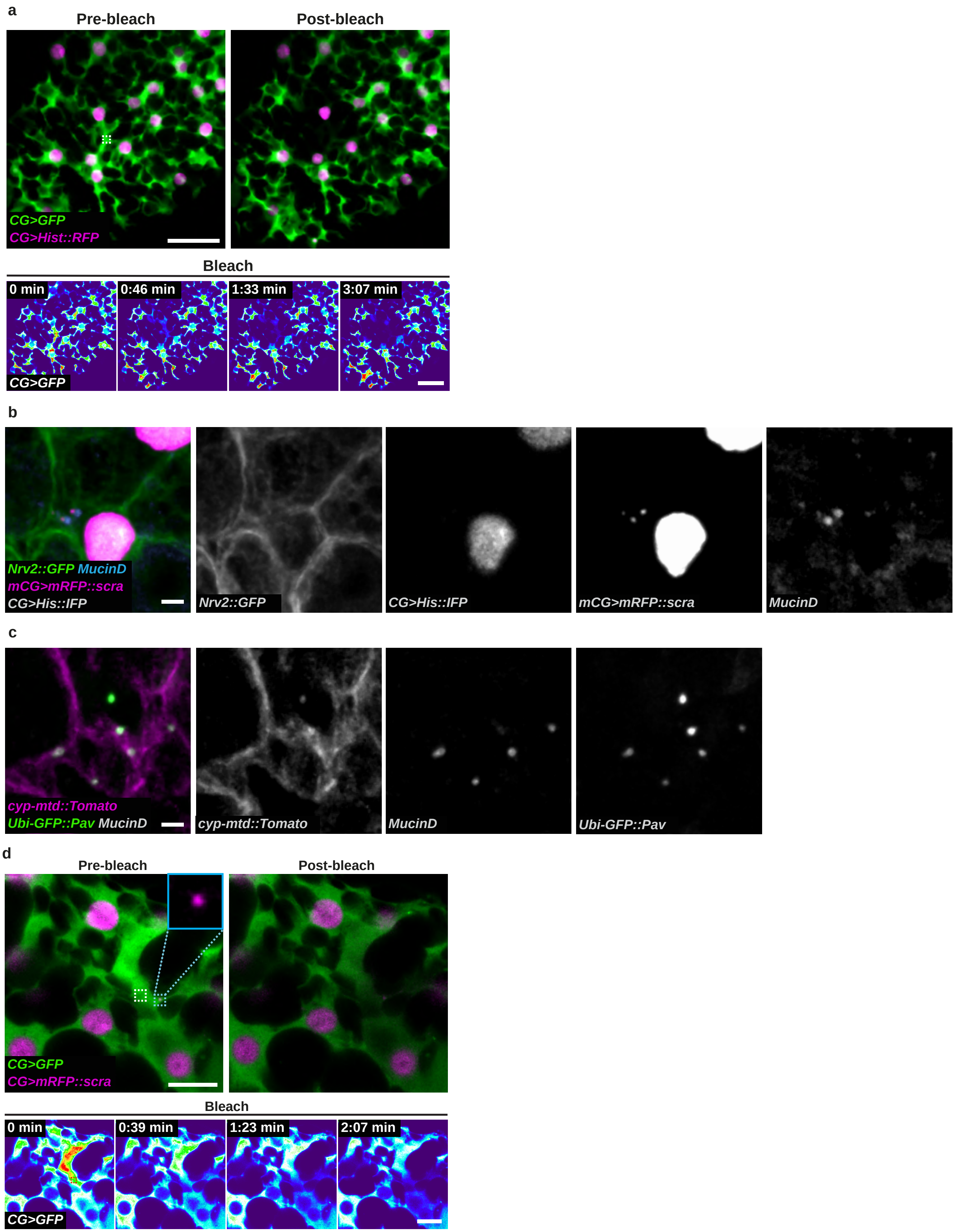

Rujano et al. Figure 4. Supplement 1

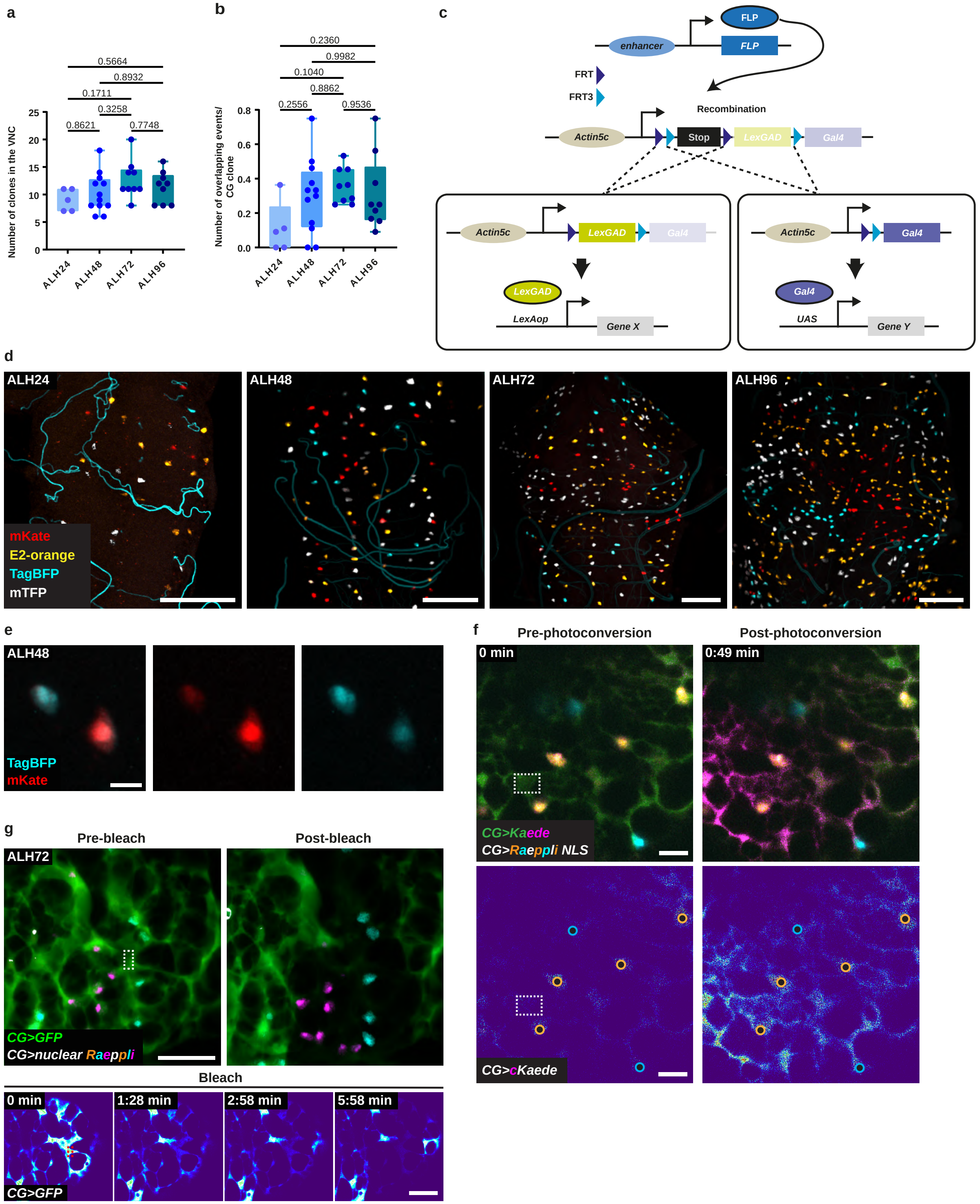

### Rujano et al. Figure 5. Supplement 1

**a**

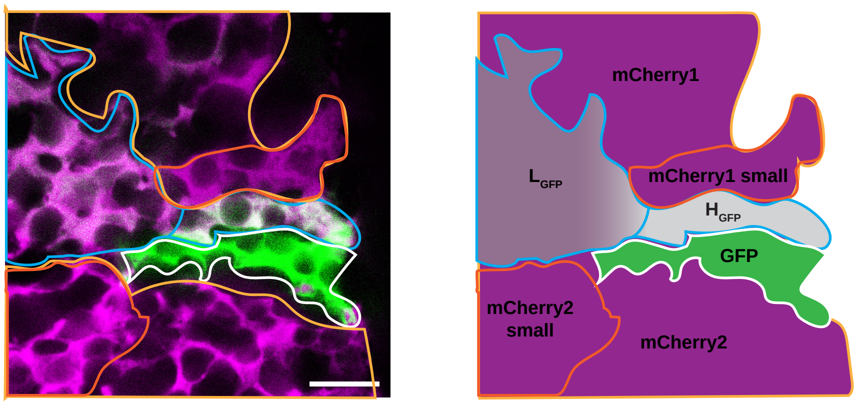


**b**

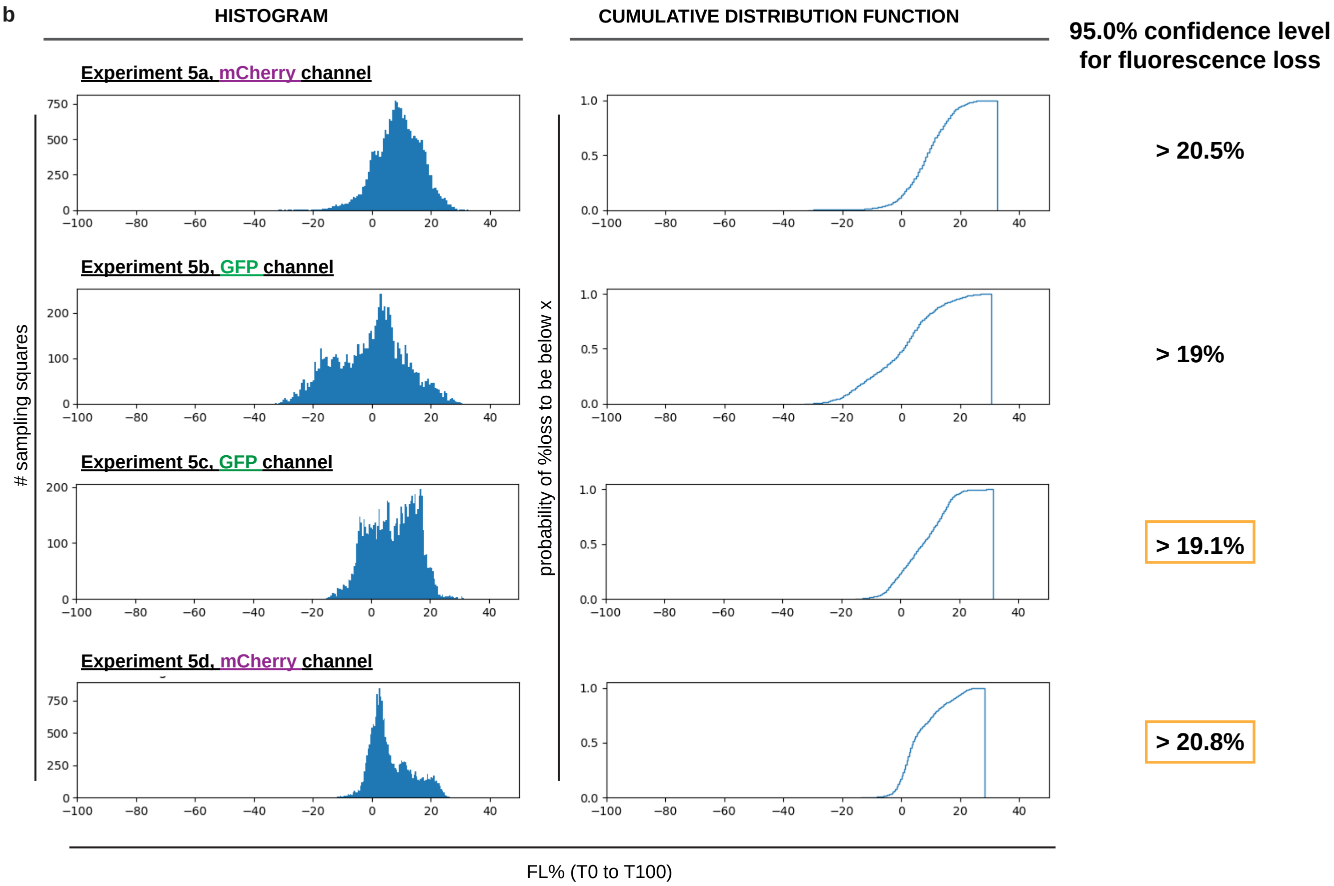


**c**

| | Zone | GFP | mCherry1 | mCherry1 small | mCherry 2 | mCherry 2 small | $H_{GFP}$ | | $L_{GFP}$ | |
|---|---|---|---|---|---|---|---|---|---|---|
| | Fluorophore | GFP | mCherry | mCherry | mCherry | mCherry | GFP | mCherry | GFP | mCherry |
| 5a | IMEAN T0 | 132.02 | 54.43 | 70.54 | 68.86 | 77.77 | 88.13 | 80.27 | 17.46 | 68.91 |
| | IMEAN T100 | 17.7 | 53.04 | 66.44 | 60.65 | 69.76 | 48.85 | 74.81 | 15.36 | 61.09 |
| | % FL | 86.59 | 2.55 | 5.81 | 11.92 | 10.30 | 44.57 | 6.80 | 12.03 | 11.35 |
| 5b | IMEAN T0 | 18.98 | 60.99 | 71.69 | 56.02 | 71.19 | 41.84 | 73.58 | 16.79 | 63.63 |
| | IMEAN T100 | 15.64 | 52.79 | 57.71 | 55.23 | 70.72 | 35.94 | 15.87 | 17.07 | 63.35 |
| | % FL | 17.60 | 13.44 | 19.50 | 1.41 | 0.66 | 14.10 | 78.43 | -1.67 | 0.44 |
| 5c | IMEAN T0 | 19.22 | 50.81 | 55.13 | 56.78 | 71.83 | 44.96 | 27.56 | 19.95 | 74.01 |
| | IMEAN T100 | 16.23 | 47.96 | 51.31 | 46.46 | 56.2 | 40.16 | 25.29 | 19.32 | 22.2 |
| | % FL | 15.56 | 5.61 | 6.93 | 18.18 | 21.76 | 10.68 | 8.24 | 3.16 | 70.00 |
| 5d | IMEAN T0 | 18.85 | 53.19 | 51.28 | 47.57 | 56.06 | 43.25 | 29.5 | 20.06 | 25.3 |
| | IMEAN T100 | 12.97 | 50.02 | 49.62 | 47.02 | 55.43 | 39.44 | 27.77 | 7.3 | 21.66 |
| | % FL | 31.19 | 5.96 | 3.24 | 1.16 | 1.12 | 8.81 | 5.86 | 63.61 | 14.39 |

Rujano et al. Figure 5. Supplement 2

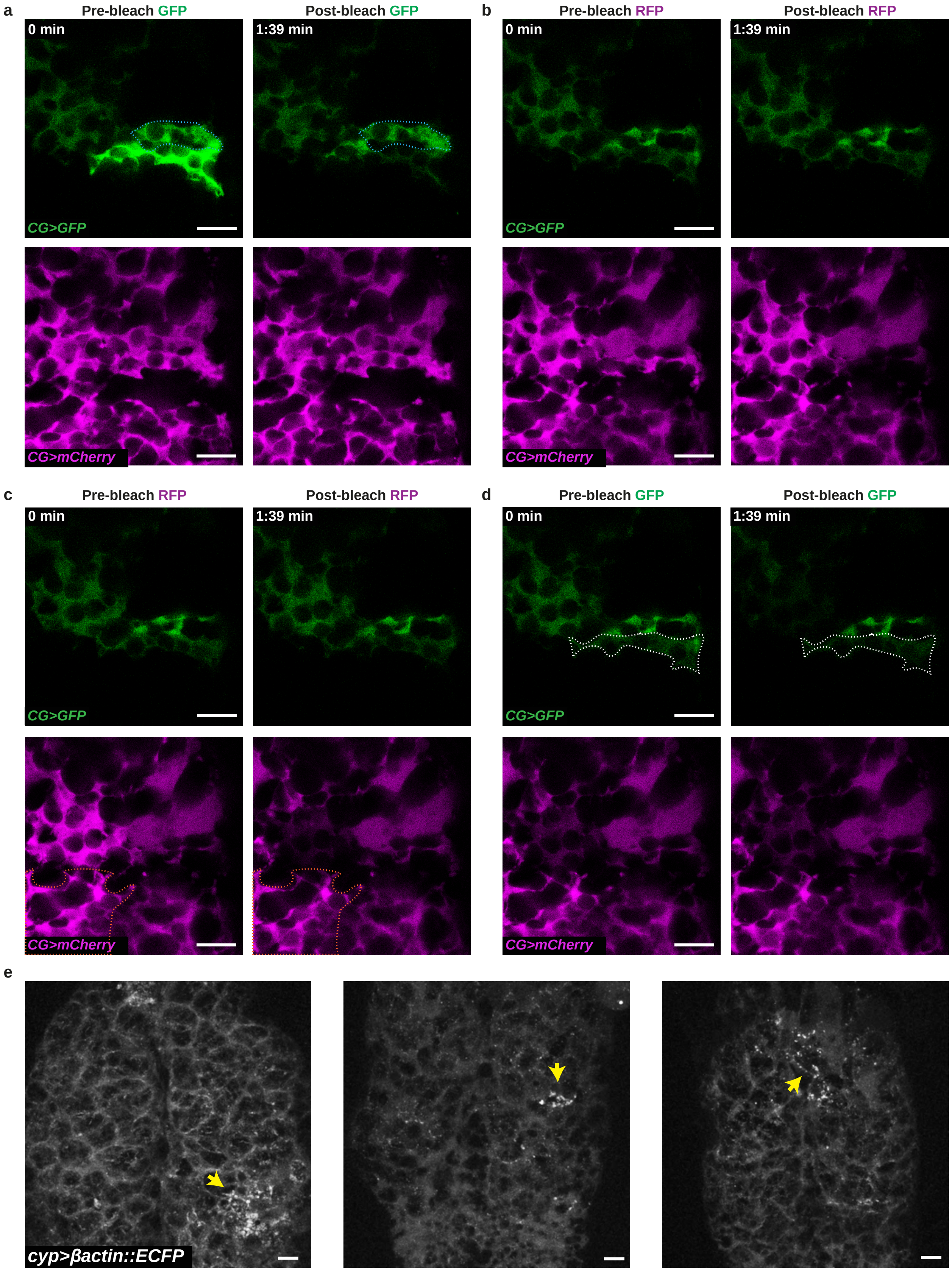

Rujano et al. Figure 6. Supplement 1

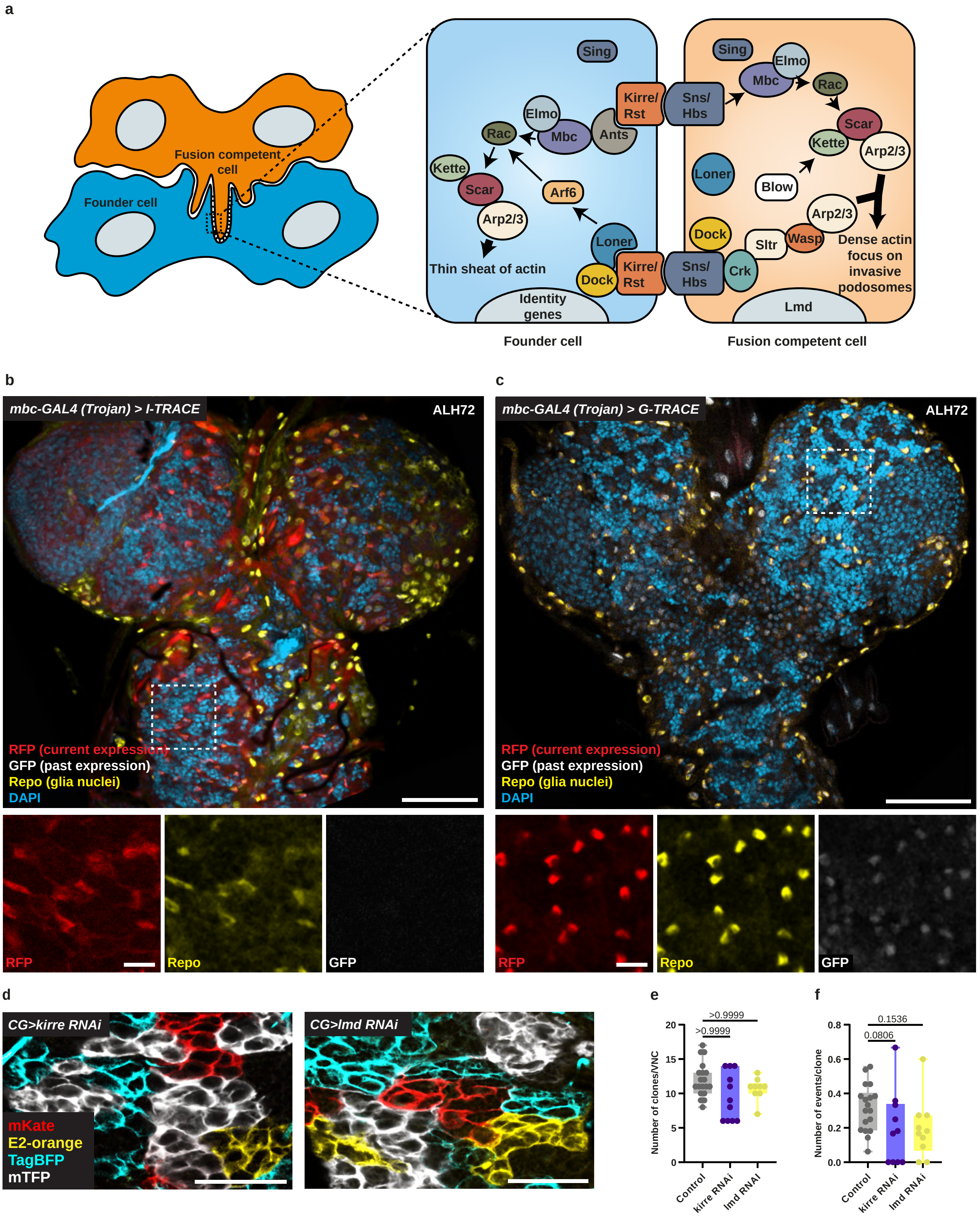

Rujano et al. Figure 7. Supplement 1

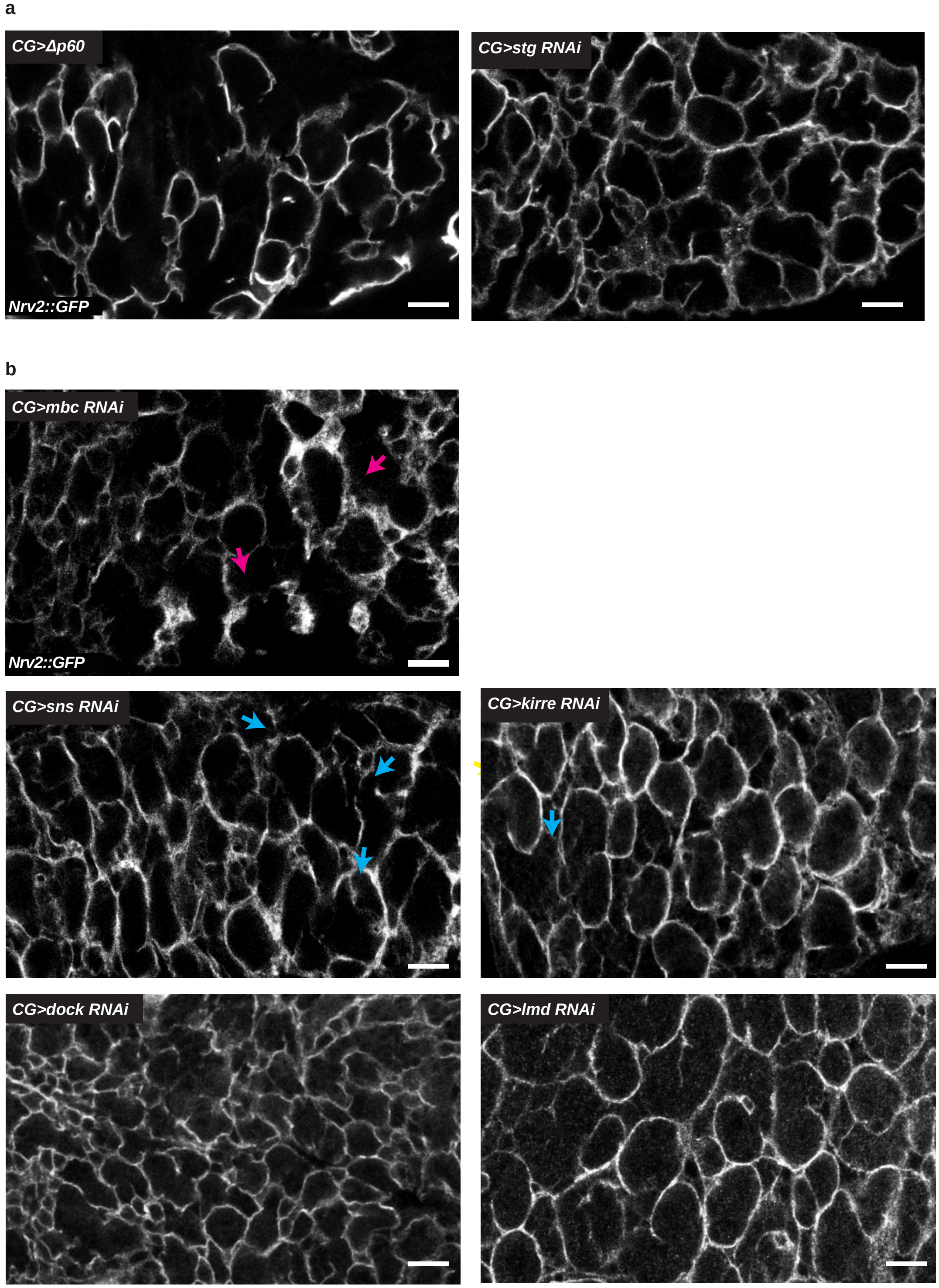
